# Quantitative relations among causality measures with applications to pulse-output nonlinear network reconstruction

**DOI:** 10.1101/2023.04.02.535284

**Authors:** Zhong-qi K. Tian, Kai Chen, Songting Li, David W. McLaughlin, Douglas Zhou

## Abstract

The causal connectivity of a network is often inferred to understand the network function. It is arguably acknowledged that the inferred causal connectivity relies on the causality measure one applies, and it may differ from the network’s underlying structural connectivity. However, the interpretation of causal connectivity remains to be fully clarified, in particular, how causal connectivity depends on causality measures and how causal connectivity relates to structural connectivity. Here, we focus on nonlinear networks with pulse signals as measured output, *e*.*g*., neural networks with spike output, and address the above issues based on four intensively utilized causality measures, *i*.*e*., time-delayed correlation coefficient, time-delayed mutual information, Granger causality, and transfer entropy. We theoretically show how these causality measures are related to one another when applied to pulse signals. Taking the simulated Hodgkin-Huxley neural network and the real mouse brain network as two illustrative examples, we further verify the quantitative relations among the four causality measures and demonstrate that the causal connectivity inferred by any of the four well coincides with the underlying network structural connectivity, therefore establishing a direct link between the causal and structural connectivity. We stress that the structural connectivity of networks can be reconstructed pairwise without conditioning on the global information of all other nodes in a network, thus circumventing the curse of dimensionality. Our framework provides a practical and effective approach for pulse-output network reconstruction.

**Significance Statement:** Inferring network connectivity is a key challenge in many diverse scientific fields. We investigate networks with pulse signal as measured output and solve the above reverse-engineering issue by establishing a direct link between the network’s causal connectivity and structural connectivity. Here, the causal connectivity can be inferred by any one of the four causality measures, *i*.*e*., time-delayed correlation coefficient, time-delayed mutual information, Granger causality, and transfer entropy. We analytically reveal the relationship among these four measures and show that they are equally effective to fully reconstruct the network connectivity pairwise. Our work provides a practical framework to reconstruct the structural connectivity in general pulse-output nonlinear networks or subnetworks.

---

The structural connectivity of a network, such as a cortical network, is of great importance in understanding the cooperation and competition among nodes in the network Mišić and Sporns (2016); McLaughlin et al. (2000); Zhou et al. (2013a). However, it is often difficult to measure directly the structural connectivity of a network. On the other hand, with the development of experimental techniques in neuroscience, it has become feasible to record simultaneously, with high temporal resolution, the activities of many nodes in a network Stringer et al. (2019); Wheeler and Nam (2011); Stosiek et al. (2003). This provides a possibility to reveal the underlying connectivity of the network by analyzing the nodes’ activity data and identifying the causal interactions (connectivity) among them Seth (2008); Honey et al. (2009; 2010); Suárez et al. (2020); Chen et al. (2022).

There are difficulties in measuring these causal interactions: One of the most widely used statistical indicator for interaction identification is the correlation coefficient Benesty et al. (2009); Eisen et al. (1998); Ito et al. (2011) that characterizes linear dependence between two nodes. The correlation coefficient is symmetric, and thus cannot distinguish the driver-recipient relation to recover the causal connectivity Stevenson et al. (2008). To solve this, the time-delayed correlation coefficient (TDCC) Bedenbaugh and Gerstein (1997); Ito et al. (2011) was introduced to detect the direction of causal connectivity. However, TDCC, as a linear measure, may fail to capture causal interactions in nonlinear networks. As a nonlinear model-free generalization of TDCC, time-delayed mutual information (TDMI) Vastano and Swinney (1988); Schreiber (2000); Frenzel and Pompe (2007) was proposed to measure the flow of information in nonlinear networks. Despite their mathematical simplicity and computational efficiency, TDCC and TDMI cannot exclude the historical effect of signals and may encounter the issue of over estimation Frenzel and Pompe (2007); Schreiber (2000); McLaughlin et al. (2004). Granger causality (GC) Granger (1969); Bressler and Seth (2011); Barnett et al. (2018) and transfer entropy (TE) Schreiber (2000); Bossomaier et al. (2016); Borge-Holthoefer et al. (2016) were two other measures that were introduced to detect causal connectivity with the exclusion of the signal’s own historical effects. GC is based on linear regression models that assumes the causal relation can be revealed by analyzing low-order statistics of signals (up to the variance). Consequently, the validity of GC for nonlinear networks is in general questionable Li et al. (2018). In contrast, TE is a nonparametric information-theoretic measure that quantifies the causal interactions with no assumption of interaction models. However, it requires the estimation of the probability distribution of dynamical variables conditioned on the historical information in networks, which makes TE suffer from the curse of dimensionality in practical applications to network systems with many nodes Runge et al. (2012); Newell and Cheng (2016); Bach (2017). It is known that the causal connectivity inferred by different causality measures can be inconsistent with each other Marbach et al. (2010); De Smet and Marchal (2010); Zou et al. (2009). With one exception [TE has been proven to be equivalent to GC for Gaussian variables Barnett et al. (2009)], there is little understanding of the relationships between these measurement techniques, including when they accurately (or inaccurately) predict causal interaction (connectivity).

The determination of structural connectivity from causal interactions faces even more fundamental difficulties: First, causal interactions may not be the result of direct structural connections between the nodes of the network, but rather the result of correlations arising from common external stimuli, or correlations arising from indirect connections (*X* → *Y, Y* → *Z*; resulting in correlations between *X* and *Z*, yet without direct structural connection between *X* and *Z*). Second, causal connectivity, as inferred from any of the above four measures, is statistical rather than structural Koch et al. (2002); Seth (2005); Schiele et al. (2013); Zhou et al. (2013b; 2014), i.e., causal connectivity quantifies direct statistical correlation among network nodes, whereas structural connectivity corresponds to physical connections among network nodes. In general, it is not clear if structural connectivity can be reconstructed from causal connectivity.

In this work we resolve the above issues for an important class of nonlinear networks, which we term *pulse-output networks* and which include *spiking neural networks*. The activity of each node in a pulse-output network can be described as a stochastic binary time series for the presence/absence of a pulse (spike) in each time window, e.g., spike train. Under this description, we illustrate mathematically that the four causal connectivity measures (TDCC, TDMI, GC, and TE) can be represented by one another, to their leading order in a perturbation expansion. Next, both by simulations of a Hodgkin-Huxley (HH) neural network and with experimentally measured data from mouse cortical network Allen Institute for Brain Science (2016), we verify that the mathematical relations among the four causality measures remain valid for representative samples of pulse-output networks. More importantly, for pulse-output networks, we demonstrate that the underlying structural connectivity can be recovered from the causal connectivity, itself inferred from any one of four causality measures. For pulse-output networks of both simulated Hodgkin-Huxley neural networks and a real mouse cortical network Allen Institute for Brain Science (2016), we show that the potential problems with the recovery of structural connectivity from causal connectivity, e.g., confounders and hidden nodes, can be resolved.

We emphasize that the *pulse-output* nature of a spiking neuronal network allows one to represent the neuronal signal as a binary time series, with random spike times. With the utility of this stochastic binary representation, our analytical framework shows, using only pairwise information between neurons, (i) the establishment of mathematical relationships between four common measures of causal connectivity, and (ii) the accurate pairwise predictions of structural connectivity from causal connectivity, without conditioning on the global information from all other nodes. Thus, the reconstruction circumvents the curse of dimensionality and can be applied to the reconstruction of structural connectivity in large-scale pulse-output nonlinear systems or subsystems.

## Results

### Concepts of generalized pairwise TDCC, TDMI, GC, and TE

Consider a nonlinear network of *N* nodes with dynamics given by

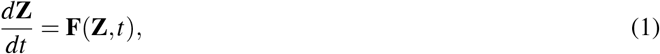

where **Z** = (*Z*_1_, *Z*_2_, …, *Z*_*N*_). We focus on the application of TDCC, TDMI, GC, and TE to each pair of nodes without conditioning on the rest of nodes in the network, accounting for the practical constraint that conditional causality measures in general require the information of the whole network that is often difficult to observe. For the ease of illustration, we denote a pair of nodes as *X* = *Z*_*i*_ and *Y* = *Z*_*j*_, and their measured time series as {*x*_*n*_} and {*y*_*n*_}, respectively.

TDCC Bedenbaugh and Gerstein (1997); Ito et al. (2011), as a function of time delay *m*, is defined by

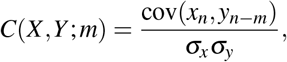

where “cov” represents the covariance, *σ*_*x*_ and *σ*_*y*_ are the standard deviations of {*x*_*n*_} and {*y*_*n*_}, respectively. A positive (negative) value of *m* indicates the calculation of causal value from *Y* to *X* (from *X* to *Y*), and nonzero *C*(*X,Y* ; *m*) indicates the existence of causal interaction between *X* and *Y*. Without loss of generality, we consider the case of positive *m* in the following discussions, that is, the causality measure from *Y* to *X*.

In contrast to the linear measure TDCC, TDMI is a model-free method being able to characterize nonlinear causal interactions Vastano and Swinney (1988); Schreiber (2000); Frenzel and Pompe (2007). TDMI from *Y* to *X* is defined by

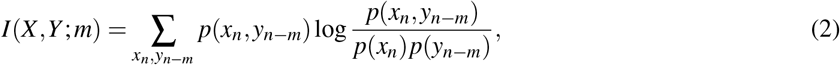

where *p*(*x*_*n*_, *y*_*n*−*m*_) is the joint probability distribution of *x*_*n*_ and *y*_*n*−*m*_, *p*(*x*_*n*_) and *p*(*y*_*n*−*m*_) are the corresponding marginal probability distributions. *I*(*X,Y* ; *m*) is non-negative and vanishes if and only if *x*_*n*_ and *y*_*n*−*m*_ are independent Frenzel and Pompe (2007). Nonzero *I*(*X,Y* ; *m*) implies the existence of causal interaction from *Y* to *X* for a positive *m*.

It has been noted that TDCC and TDMI could overestimate the causal interactions when a signal has a long memory Frenzel and Pompe (2007); Schreiber (2000); McLaughlin et al. (2004). As an alternative, GC was proposed to overcome the issue of overestimation based on linear regression Granger (1969); Guo et al. (2008); Bressler and Seth (2011). The auto-regression for *X* is represented by 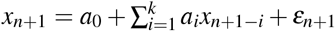, where {*a*_*i*_} are the *estimated* auto-regression coefficients and *ε*_*n*+1_ is the residual. By including the historical information of *Y* with a message length *l* and a time-delay *m*, the joint regression for *X* is represented by 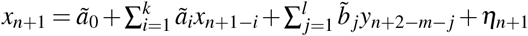, where {*ã*_*i*_} and 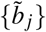 are the *estimated* joint regression coefficients, and *η*_*n*+1_ is the corresponding residual. If there exists a causal interaction from *Y* to *X*, then the prediction of *X* using the linear regression models shall be improved by additionally incorporating the historical information of *Y*. Accordingly, the variance of residual *η*_*n*+1_ is smaller than that of *ε*_*n*+1_. Based on this concept, the GC value from *Y* to *X* is defined by

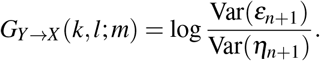

The GC value is also non-negative and vanishes if and only if 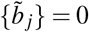, *i*.*e*., the variance of residual *ε*_*n*+1_ for *X* cannot be reduced by including the historical information of *Y*. Note that, by introducing the time-delay parameter *m*, the GC analysis defined above generalizes the conventional GC analysis, as the latter corresponds to the special case of *m* = 1.

GC assumes that the causal interaction can be fully captured by the variance reduction in the linear regression models, which is valid for Gaussian signals but not for more general signals. As a nonlinear extension of GC, TE was proposed to describe the causal interaction from the information theoretic perspective Schreiber (2000). The TE value from *Y* to *X* is defined by

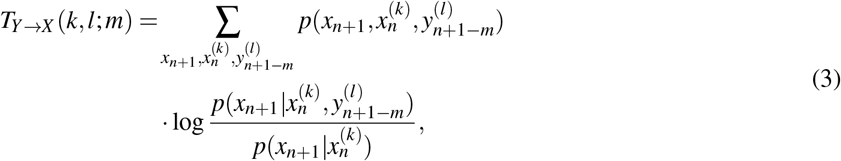

where the shorthand notation 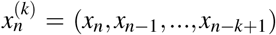 and 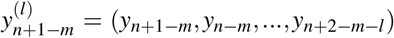, *k,l* indicate the length (order) of historical information of *X* and *Y*, respectively. Similar to GC, the time-delay parameter *m* is introduced that generalizes the conventional TE, the latter of which corresponds to the case of *m* = 1. TE is non-negative and vanishes if and only if 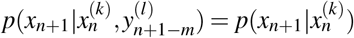, *i*.*e*., the uncertainty of *x*_*n*+1_ is not affected regardless of whether the historical information of *Y* is taken into account.

In this work, we investigate the mathematical relations among TDCC, TDMI, GC, and TE by focusing on nonlinear networks described by Eq. 1 with pulse signals as measured output, *e*.*g*., the spike trains measured in neural networks.

Consider a pair of nodes *X* and *Y* in the network of *N* nodes, and denote their pulse-output signals by

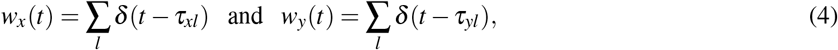

respectively, where *δ* (·) is the Dirac delta function, and {*τ*_*xl*_} and {*τ*_*yl*_} are the output time sequences of nodes *X* and *Y* determined by Eq. 1, respectively. With the sampling resolution of Δ*t*, the pulse-output signals are measured as binary time series {*x*_*n*_} and {*y*_*n*_}, where *x*_*n*_ = 1 (*y*_*n*_ = 1) if there is a pulse signal, *e*.*g*., a spike generated by a neuron, of *X* (*Y*) occurred in the time window [*t*_*n*_, *t*_*n*_ + Δ*t*), and *x*_*n*_ = 0 (*y*_*n*_ = 0) otherwise, *i*.*e*.,

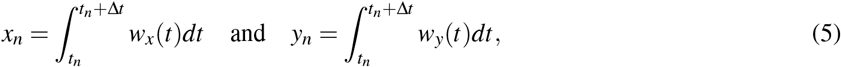

and *t*_*n*_ = *n*Δ*t*. Note that the value of Δ*t* is often chosen to make sure that there is at most one pulse signal in a single small enough time window. In the stationary state, the responses *x*_*n*_ and *y*_*n*_ can be viewed as stochastic processes when the network is driven by stochastic external inputs. In such a case, for the sake of simplicity, we denote *p*_*x*_ = *p*(*x*_*n*_ = 1), and define 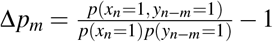, measuring the dependence between *x*_n_ and *y*_*n*−*m*_.

### Mathematical relation between TDMI and TDCC

For the relation between TDCC and TDMI when applied to nonlinear networks with pulse-output signals, we prove the following theorem:

#### Theorem 1.

*For nodes X and Y with pulse-output signals given in Eqs. 4 and 5, we have*

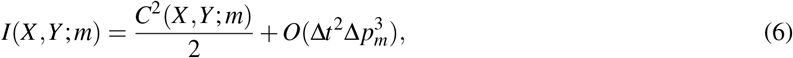

*where the symbol* “*O*” *stands for the order*.

*Proof*. The basic idea is to Taylor expand TDMI in Eq. 2 with respect to the term 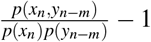 (the detailed derivation can be found in *SI Appendix, Supporting Information Text 1B*), then we arrive at the following expression:

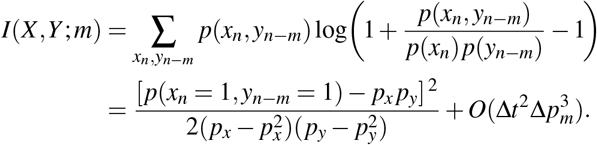

Since TDCC can be written as

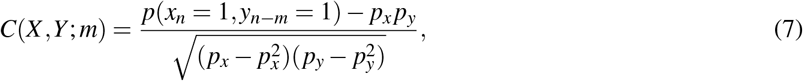

we have

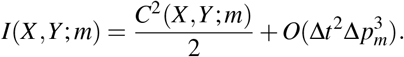

### Mathematical relation between GC and TDCC

We next derive the relation between GC and TDCC as follows:

#### Theorem 2.

*For nodes X and Y with pulse-output signals given in Eqs. 4 and 5, we have*

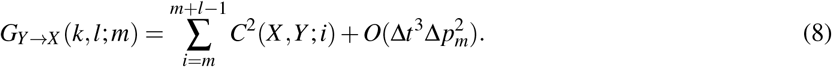

*Proof*. From the definition, GC can be represented by the covariances of the signals Barnett et al. (2009) as

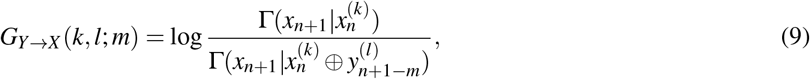

where G(**x**|**y**) = cov(**x**) − cov(**x, y**)cov(**y**)^−1^cov(**x, y**)^*T*^ for random vectors **x** and **y**, cov(**x**) and cov(**y**) denote the covariance matrix of **x** and **y**, respectively, and cov(**x, y**) denote the cross-covariance matrix between **x** and **y**. The symbol *T* is the transpose operator and ⊕ denotes the concatenation of vectors.

We first prove that the auto-correlation function (ACF) of binary time series {*x*_*n*_} as a function of time delay takes the order of Δ*t* (see *SI Appendix, Supporting Information Text 1C* for the proof, Fig. S1). Accordingly, we have

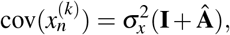

where **Â** = (*â*_*i j*_), *â*_*i j*_ = *O*(Δ*t*), and **I** is the identity matrix. Hence,

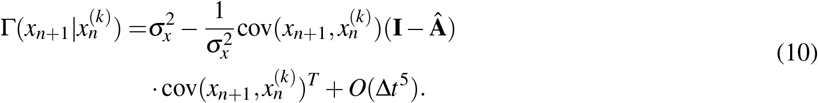

In the same way, we have

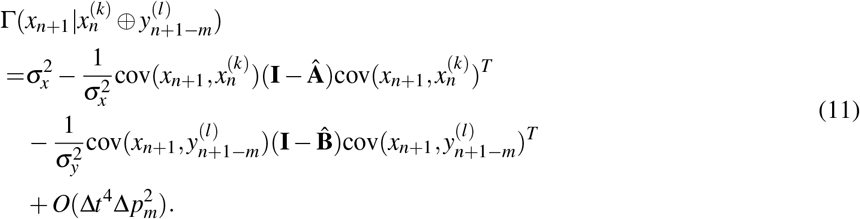

where 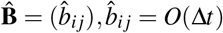. Substituting Eqs. 10 and 11 into Eq. 9 and Taylor expanding Eq. 9 with respect to Δ*t*, we can obtain

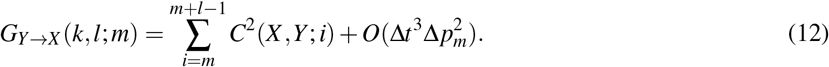

The detailed derivation of Eqs. 10, 11, and 12 can be found in *SI Appendix, Supporting Information Text 1C*.

### Mathematical relation between TE and TDMI

From the definitions of TE and TDMI, TE can be regarded as a generalization of TDMI conditioning on the signals’ historical information additionally. To rigorously establish their relationship, we require that 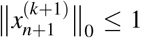 and 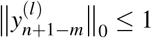 in the definition of TE given in Eq. 3, where ‖· ‖_0_ denotes the *l*_0_ norm of a vector, *i*.*e*., the number of nonzero elements in a vector. This assumption indicates that the length of historical information used in the TE framework is shorter than the minimal time interval between two consecutive pulse-output signals. With this condition, we mathematically establish the following theorem:

#### Theorem 3.

*For nodes X and Y with pulse-output signals given in Eqs. 4 and 5, under the assumption that* 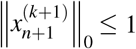 *and* 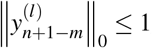, *we have*

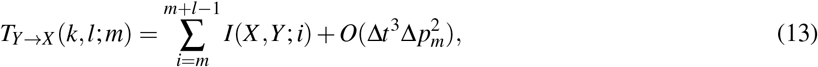

*where T*_*Y*→*X*_ *is defined in Eq. 3*.

*Proof*. To simplify the notation, we denote 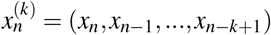 and 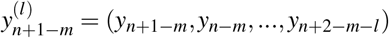 as *x*^−^ and *y*^−^, respectively. From Eq. 3, we have

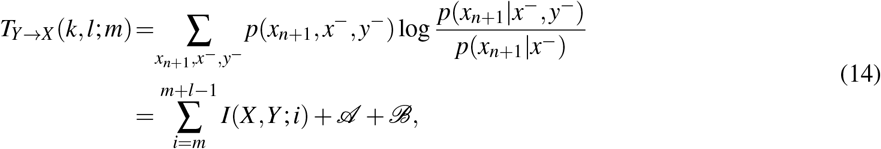

where

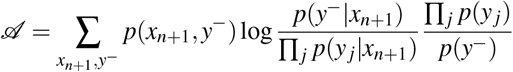

and

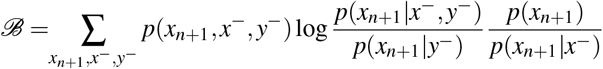

and ∏ _*j*_ in *𝒜* represents 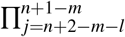. The detailed derivation of Eq. 14 can be found in *SI Appendix, Supporting Information Text 1D. 𝒜* and *ℬ* in Eq. 14 consist of multiple terms and the leading order of each term can be analytically calculated. For the sake of illustration, we derive the leading order of one of these terms and the leading. Without loss of generality, we assumed order of the rest terms can be estimated in a similar way. Under the assumption that 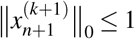 and 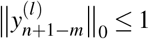, the number of nonzero components is at most one in 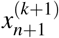 and 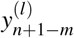. Without loss of generality, we assumed *x*_*n*+1_ = 1 and *y*_*n*+1−*m*_ = 1. In such a case, we can obtain the following expression in *𝒜*,

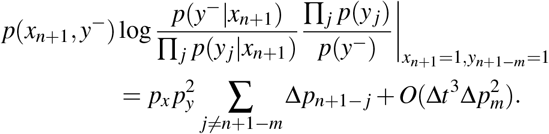

We can further show that the leading order of all the terms in *𝒜* cancel each other out (*SI Appendix, Supporting Information Text 1D*), thus we have 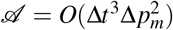. Similarly, we can also show *ℬ* = *O*(Δ*t*^3^Δ*p*^2^) (*SI Appendix, Supporting Information Text 1D*), and thus

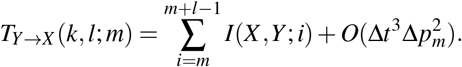

### Mathematical relation between GC and TE

From Theorems 1-3, we can prove the following theorem (see details in *SI Appendix, Supporting Information Text 1E*):

#### Theorem 4.

*For nodes X and Y with pulse-output signals given in Eqs. 4 and 5, under the assumption that* 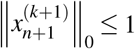 *and* 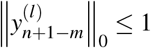, *we have*

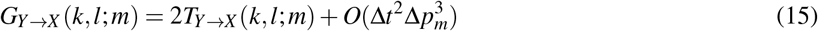

*where T*_*Y*→*X*_ *is defined in Eq. 3*.

Note that in Theorems 3 and 4 we require the condition 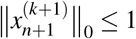 and 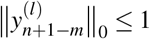 to establish the relations of causality measures. However, the relations of the four causality measure remain to hold approximately in the absence of this condition, as will be discussed below. We summarize the relations among the four causality measures in Fig. 1.

**Figure 1:**
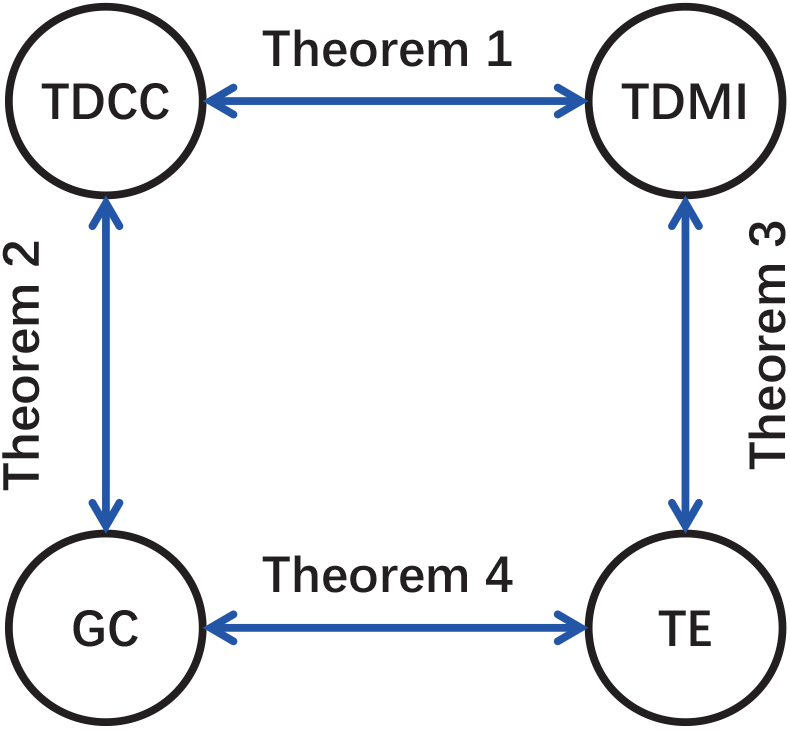
Mathematical relations among TDCC, TDMI, GC, and TE estiblished by four theorems.

### Mathematical relations of causality measures verified in HH neural networks

To verify the relations among the causality measures derived above, as an illustrative example, we apply generalized pairwise TDCC, TDMI, GC, and TE to the HH neural network described in *Materials and Methods*. We first consider a pair of neurons denoted by *X* and *Y* with unidirectional connection from *Y* to *X* in an HH network containing 10 excitatory neurons driven by homogeneous Poisson inputs. Let {*τ*_*xl*_} and {*τ*_*yl*_} be the ordered spike times of neuron *X* and *Y* in the HH network respectively and denote their spike trains as *w*_*x*_(*t*) = ∑_*l*_ *δ* (*t* − *τ*_*xl*_) and *w*_*y*_(*t*) = ∑_*l*_ *δ* (*t* − *τ*_*yl*_), respectively. With a sampling resolution of Δ*t*, the spike train is measured as a binary time series as described above. To numerically verify the above theorems, we then check the order of the remainders in Eqs. 6, 8, 13, and 15 in terms of Δ*t* and Δ*p*_*m*_. Note that Δ*p*_*m*_, the measure of the dependence between *X* and *Y*, is insensitive to sampling resolution Δ*t* (*SI Appendix*, Fig. S2). Therefore, by varying sampling interval Δ*t* and coupling strength *S* (linearly related to Δ*p*_*m*_) respectively, the orders of the remainders are consistent with those derived in Eqs. 6, 8, 13, and 15 as shown in Fig. 2. In addition, Fig. 3 verifies the relations among the causality measures by changing other parameters. For example, in Eqs. 8, 13, and 15, the four causality measures are proved to be independent of the historical length *k*, which is numerically verified in Fig. 3*A*. And although the values of GC and TE rely on the historical length *l* in Fig. 3*B*, the mathematical relations among the four causality measures revealed by Theorems 1-4 hold for a wide range of *l*. Recall that, in order to establish rigorously the relations of the causality measures of Theorems 3 and 4, our proofs required the assumption 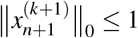 and 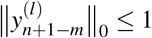. This assumption can often be satisfied: For example, in the above simulations, the memory time of the neuron is about 20 ms, while the inter-spike interval is around 100 ms. However, even when the assumption breaks down in the regime of high firing rate, the mathematical relations among the four causality measures remain to hold approximately, as examined numerically in *SI Appendix*, Fig. S3.

**Figure 2:**
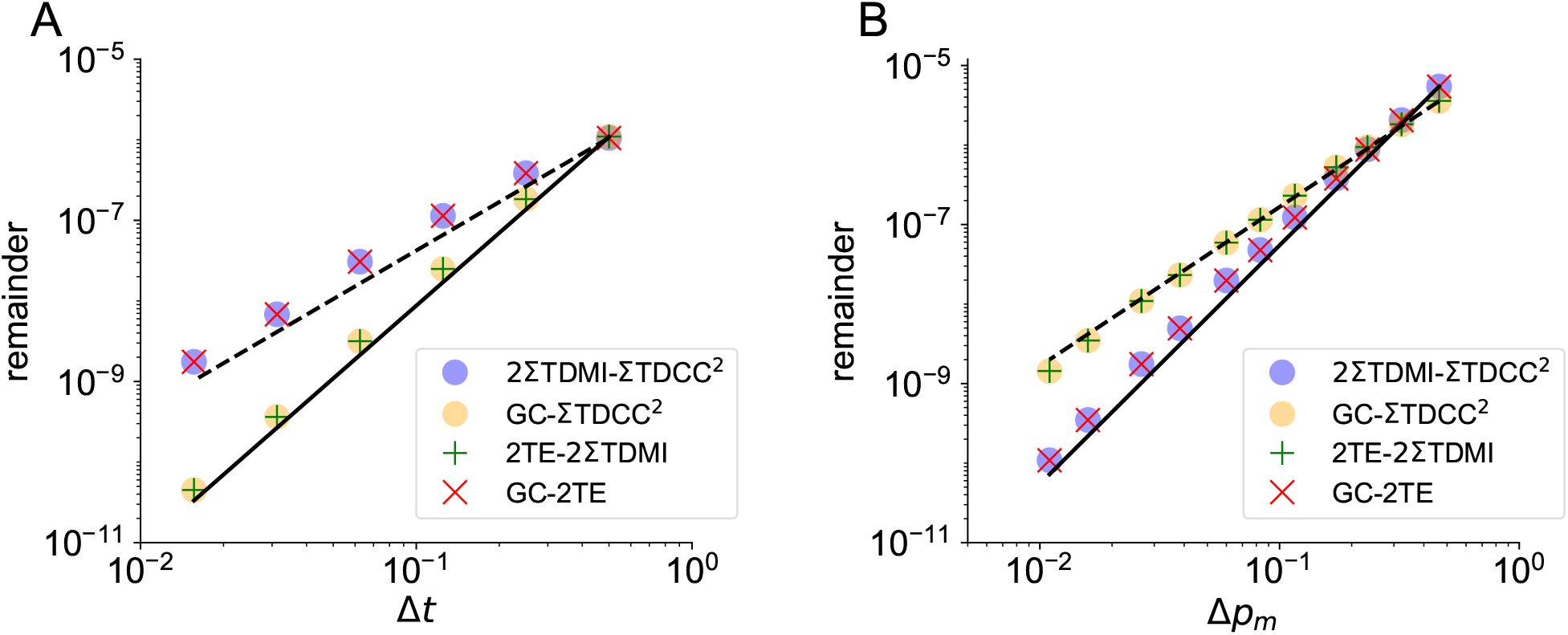
Numerical verification of the convergence order of the remainders in terms of (*A*) Δ*t* and (*B*) Δ*p*_*m*_. The convergence orders for Δ*t* in (*A*) and Δ*p*_*m*_ in (*B*) agree well with Theorems 1-4 (*R*^2^ *>* 0.998). The gray dashed and solid lines indicate the 2^nd^-order and 3^rd^-order convergence, respectively. The four causal measures are calculated from a pair of unidirectionally connected neurons in an HH network of 10 excitatory neurons randomly connected with probability 0.25. The parameters are set as *k* = *l* = 5 and *m* = 6 (time delay is 3 ms), *S* = 0.02 mS·cm^−2^ in (*A*), and Δ*t* = 0.5 ms in (*B*).

**Figure 3:**
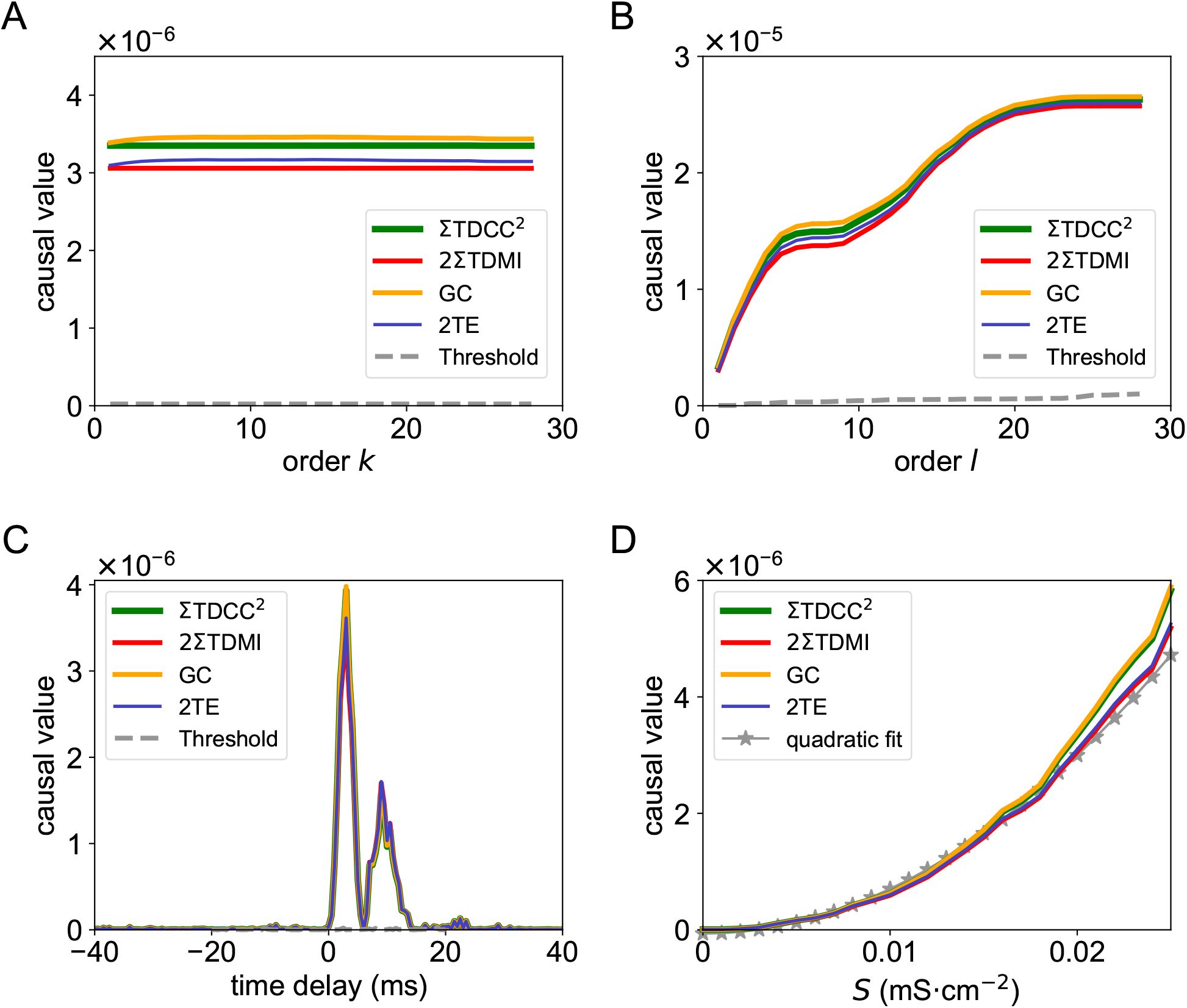
Dependence of causal values on parameters of (*A*) order *k*, (*B*) order *l*, (*C*) time delay, and (*D*) coupling strength *S* obtained from the same pair of neurons in the HH network in Fig. 2. In (*C*), a positive (negative) time delay indicates the calculation of causal values from neuron *Y* to neuron *X* (from *X* to *Y*). The green curve represents the summation of squared TDCC *C*(*X,Y* ; *m*), the red curve represents twice of the summation of TDMI *I*(*X,Y* ; *m*), the orange curve stands for GC *G*_*Y*→*X*_ (*k, l*; *m*), and the blue curve stands for twice of TE *T*_*Y*→*X*_ (*k, l*; *m*). The curves virtually overlap in (A)-(D) (all significantly greater than those of randomly surrogate time series, *p <* 0.05). The gray dashed curve in (A)-(C) is the significance level of causality for two unconnected neurons in the same HH network. The gray star curve in (D) is the quadratic fit of causal values with respect to different *S*. The parameters are set as (*A*): *l* = 1, *S* = 0.02 mS·cm^−2^, Δ*t* = 0.5 ms, *m* = 6; (*B*): *k* = 1, *S* = 0.02 mS·cm^−2^, Δ*t* = 0.5 ms, *m* = 6; (*C*): *k* = *l* = 1, *S* = 0.02 mS·cm^−2^, Δ*t* = 0.5 ms; (*D*): *k* = *l* = 1, Δ*t* = 0.5 ms, *m* = 6.

We next verify the mathematical relations among the causality measures for the parameter of time delay *m* by fixing parameters *k* and *l*. In principle, the value of *k* and *l* in GC and TE shall be determined by the historical memory of the system. To reduce the computational cost Gourévitch and Eggermont (2007); Vicente et al. (2011); Ito et al. (2011), we take *k* = *l* = 1 for all the results below. It turns out that this parameter choice works well for pulse-output networks because of the short memory effect in general, as will be further discussed later. Fig. 3*C* shows the mathematical relations hold for a wide range of the time-delay parameter used in computing the four causality measures (see more examples in *SI Appendix*, Fig. S4).

We further examine the robustness of the mathematical relations among TDCC, TDMI, GC, and TE by scanning the parameters of the coupling strength *S* between the HH neurons and external Poisson input strength and rates. As shown in Fig. 3*D*, the values of the four causality measures with different coupling strength are very close to one another. Their relations also hold for a wide range of external Poisson input parameters (*SI Appendix*, Fig. S5). From the above, the mathematical relations among TDCC, TDMI, GC, and TE described in Theorems 1-4 are verified in the HH network.

### Relation between structural connectivity and causal connectivity in HH neural networks

We next discuss the relation between the inferred causal connectivity and the structural connectivity. Note that the causal connectivity inferred by these measures is statistical rather than structural Koch et al. (2002); Seth (2005); Schiele et al. (2013), *i*.*e*., the causal connectivity quantifies the direct statistical correlation or dependence among network nodes, whereas the structural connectivity corresponds to physical connections among network nodes. Therefore, a precise relationship between causal connectivity and structural connectivity has been unclear. In Fig. 3*C*, the peak causal value from *Y* to *X* (at time delay around 3 ms, *m* = 6) is greater than the significance level (the gray dashed line in Fig. 3*C*), while the causal value from *X* to *Y* is not. Based on this, the inferred direct causal connections between *X* and *Y* are consistent with the underlying structural connections. From now on, we adopt peak causal values, *m* = 6, to represent the causal connectivity unless noted explicitly.

To investigate the validity of this consistency in larger networks, we further investigate a larger HH network (100 excitatory neurons) with random connectivity structure and homogeneous coupling strength (*SI Appendix*, Fig. S6A). As shown in Fig. 4*A*, the distributions of all four causal values across all pairs of neurons virtually overlapped, which again verify their mathematical relations given by Theorems 1-4. In addition, as the network size increases from 10 to 100 neurons, the distributions of the causality measures in Fig. 4*A* exhibits a bimodal structure with a clear separation of orders. By mapping the causal values with the structural connectivity, we find that the right bump of the distributions with larger causal values corresponded to connected pairs of neurons, while the left bump with smaller causal values corresponded to unconnected pairs. The well separation of the two modals indicates that the underlying structural connectivity in the HH network can be accurately estimated from the causal connectivity.

**Figure 4:**
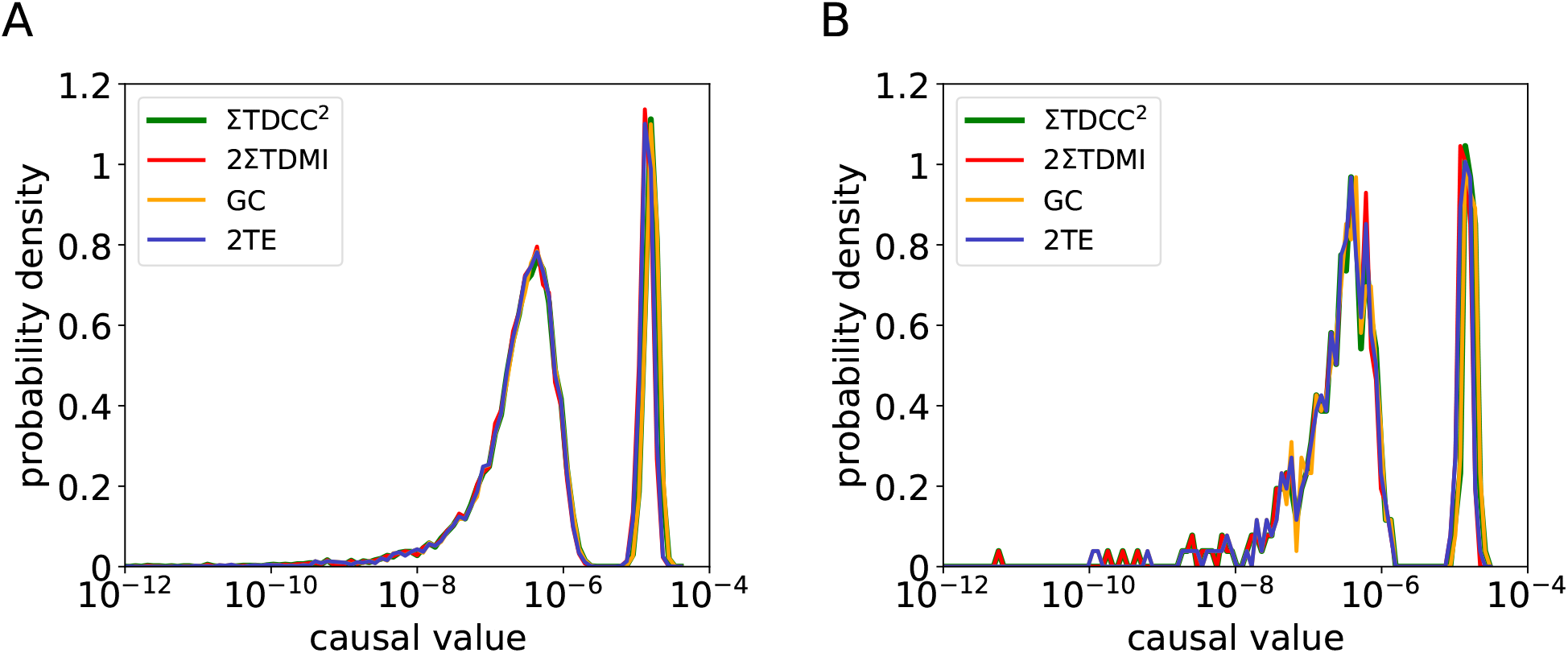
Distributions of causal values in an HH network of 100 excitatory neurons randomly connected with probability 0.25. (*A*) The distribution of causal values of each pair of neurons in the whole network. (*B*) The distribution of causal values of each pair of neurons in an HH subnetwork of 20 excitatory neurons. The parameters are set as *k* = *l* = 1, *S* = 0.02 mS·cm^−2^, Δ*t* = 0.5 ms, and *m* = 6. The colors are the same as those in Fig. 3 and the curves nearly overlap.

The performance of this reconstruction approach can be quantitatively characterized by the receiver operating characteristic (ROC) curve and the area under the ROC curve (AUC) Fawcett (2006); Marbach et al. (2010); Carter et al. (2016) (see *Materials and Methods*). It is found that the AUC value becomes 1 when applying any of these four causality measures (*SI Appendix*, Fig. S6B), which indicates that the structural connectivity of the HH network could be reconstructed with 100% accuracy. We point out that the reconstruction of network connectivity based on causality measures is achieved by calculating the causal values between each pair of neurons that requires no access to the activity data of the rest of neurons. Therefore, this inference approach can be applied to a subnetwork when the activity of neurons outside the subnetwork is not observable. For example, when a subnetwork of 20 excitatory HH neurons is observed, the structural connectivity of the subnetwork can still be accurately reconstructed without knowing the information of the rest 80 neurons in the full network as shown in Fig. 4*B*. In such a case, the AUC values corresponding to the four causality measures are 1 (*SI Appendix*, Fig. S6C). In addition, we have also shown that, for HH networks with *heterogeneous* coupling strength (for example, the coupling strength follows a log-normal distribution), accurate reconstructions of network structural connectivity can still be achieved (*SI Appendix*, Fig. S7).

### Mechanism underlying network connectivity reconstruction by causality measures

We next demonstrate the mechanism underlying the validity of pairwise inference of pulse-output signals in the reconstruction of network structural connectivity. It has been frequently noticed that pairwise causal inference may potentially fail to distinguish direct interactions from indirect ones Stevenson et al. (2008). For example, in the case that *Y* → *W* → *X* where “→” denotes a direct connection, the indirect interaction from *Y* to *X* may possibly be mis-inferred as a direct interaction via pairwise causality measures. However, pulse-output signals circumvent such spurious inferences as explained below. Here we take TDCC as an example to elucidate the underlying reason of successful reconstruction. If we denote *δ p*_*Y*→*X*_ = *p*(*x*_*n*_ = 1|*y*_*n*−*m*_ = 1) − *p*(*x*_*n*_ = 1|*y*_*n*−*m*_ = 0) as the increment of probability of generating a pulse output for neuron *X* induced by a pulse-output signal of neuron *Y* at *m* time step earlier, we have 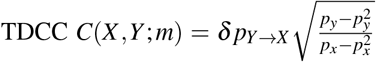 through Eq. 7. For the case of *Y* → *W* → *X*, we can derive *δ p*_*Y*→*X*_ = *O*(*δ p*_*Y*→*W*_ · *δ p*_*W*→*X*_), and further *C*(*X,Y* ; *m*) = *O* (*C*(*W,Y* ; *m*) · *C*(*X,W* ; *m*)) (see derivation details about the relations among *δ p*, Δ*p*_*m*_, and causal measures in *SI Appendix, Supporting Information Text 2*, Figs. S8A-B). Because the influence of a single pulse-output signal is often small (*e*.*g*., in the HH neural network with physiologically realistic coupling strengths, we obtain |*δ p*| *<* 0.01 from simulation data), the causal value *C*(*X,Y* ; *m*) due to the indirect interaction is significantly smaller than *C*(*W,Y* ; *m*) or *C*(*X,W* ; *m*) by direct interactions as depicted in Fig. 4*A*.

We also note that the increment *δ p* is linearly dependent on the coupling strength *S* (*SI Appendix*, Fig. S8C), thus we establish a mapping between the causal and structural connectivity, in which the causal value of TDCC is proportional to the coupling strength *S* between two neurons. The mapping between causal and structural connectivity for TDMI, GC, and TE are also established in a similar way, in which the corresponding causal values are proportional to *S*^2^ as shown in Fig. 3*D*. Therefore, the application of pairwise causality measures to pulse-output signals is able to successfully reveal the underlying structural connectivity of a network. In summary, for pulse-output networks, structural connectivity can be accurately inferred from causal connectivity in a pair-wise manner that overcomes computational issues of high dimensionality. Thus, the method is potentially applicable to experimentally measured data from large-scale biological networks or subnetworks as discussed below.

### Network connectivity reconstruction with physiological experimental data

Next, we apply all four causality measures to experimental data to address the issue of validity of their mathematical relations and reconstruction of the network structural connectivity. Here, we analyze the *in vivo* spike data recorded in the mouse cortex from Allen Brain Observatory Allen Institute for Brain Science (2016) (see *Materials and Methods*). By applying the four causality measures, we infer the causal connectivity of those cortical networks. As the underlying structural connectivity of the recorded neurons in experiments is unknown, we first detect putative connected links from the distribution of causality measures, and then follow the same procedures as previously described in the HH model case using ROC and AUC in signal detection theory Fawcett (2006); Marbach et al. (2010); Carter et al. (2016) (see *Materials and Methods*) to quantify the reconstruction performance.

Because we have demonstrated the equivalence of the four causality measures, we will use TE as a representative causality measure to demonstrate a way to detect putative connections. As we have shown above, the TE values are proportional to *S*^2^. In addition, previous experimental works observed that the coupling strength *S* follows the log-normal distribution both for intra-areal and inter-areal cortical networks in mouse and monkey brains Buzsáki and Mizuseki (2014). Thus, the distribution of TE for connected pairs should follow the log-normal distribution. In addition, we assume that the distribution of TE for unconnected pairs also obeys the log-normal distribution (see *Materials and Methods*).

To validate the above assumption of two log-normal distributions of TE values, we consider an HH network where neurons are randomly connected and the network coupling strength follows the log-normal distribution as observed in experiment Buzsáki and Mizuseki (2014). In this case, although the distribution of log TE values log_10_ *T*_*Y*→*X*_ in Fig. 5*A* for synaptically connected and unconnected pairs of neurons overlap with each other, it can be well fitted by the summation of two log-normal distributions of TE values. Importantly, the fitted distributions of TE of connected and unconnected pairs agree well with those of the true connectivity set up in simulation as shown in Fig. 5*B*. Thus, we take the log-normal fitted causality measure distributions as the ground truth of structural connectivity to evaluate the performance of network connectivity reconstruction, *e*.*g*., the AUC value is 0.997 in Fig. 5*B*. Practically, an optimal inference threshold determined as the intersection of the two fitted distribution curves Dayan and Abbott (2001) can be used for network connectivity reconstruction, which is indicated by the vertical solid line in Fig. 5*B*. In addition, the above results are robust for a variety of coupling strength distributions, including uniformly distributed and Gaussian distributed coupling strength with AUC values higher than 0.95 (*SI Appendix*, Fig. S7).

**Figure 5:**
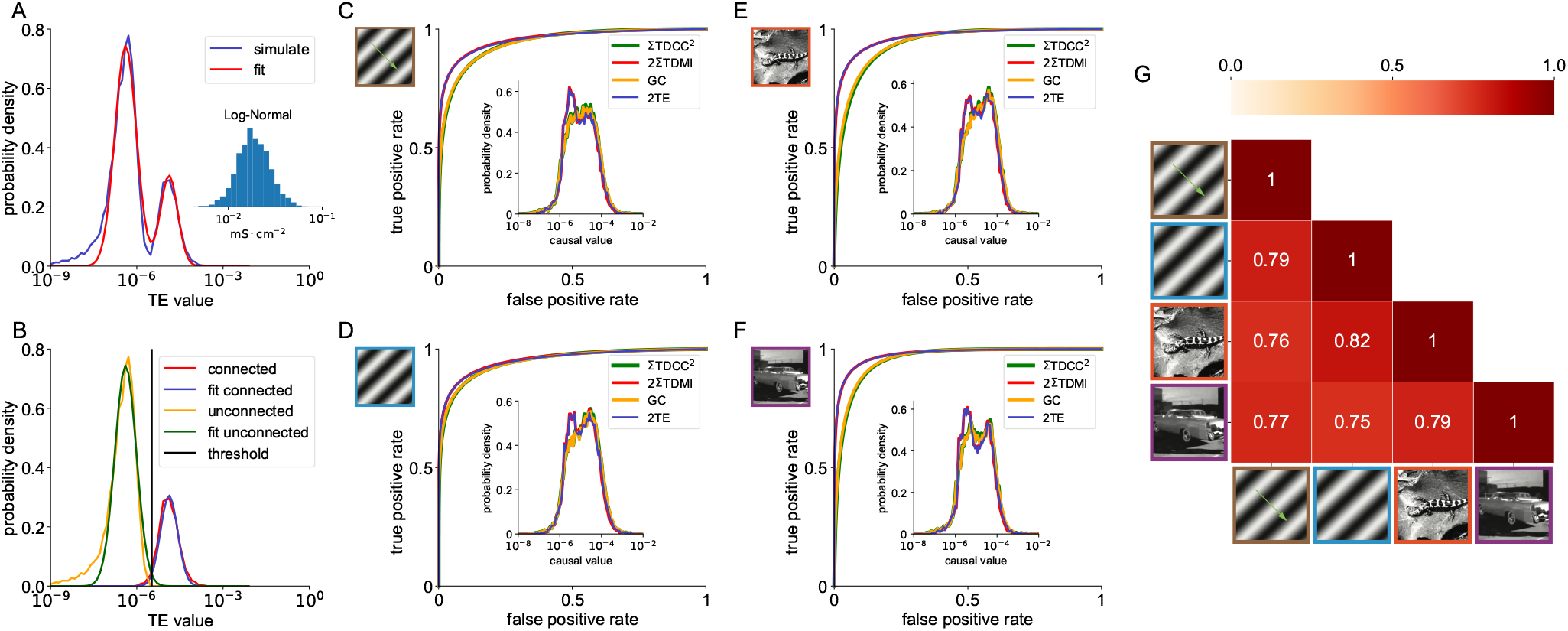
Reconstruction of structural connectivity by the assumption of mixed log-normal distribution of causal values. (*A*) Distribution of TE values in an HH network of 100 excitatory neurons. The entry *A*_*i j*_ in the adjacency matrix follows a Bernoulli distribution with probability of 0.25 being 1. For the connected pairs, *e*.*g*., *A*_*i j*_ = 1, the corresponding coupling strength from neuron *j* to neuron *i* is sampled from a log-normal distribution. The blue and red curves are the simulated and fitted distributions, respectively. Inset: histogram of structural coupling strength, *S*. (*B*) Distributions of TE values from connected and unconnected pairs of HH neurons in (*A*). The red and orange curves are the simulated TE values from connected and unconnected pairs respectively, while the blue and green curves are the fitted TE values from connected and unconnected pairs respectively which are obtained from the fitting in (*A*). The parameters are the same as those in Fig. 4. (*C-F*): ROC curves of the network composed by the recorded neurons in experiment under visual stimuli of (*C*) drifting gratings, (*D*) static gratings, (*E*) natural scenes, and (*F*) natural movie, with AUC values equal to 0.950, 0.951, 0.953, 0.967. Insets: the distribution of causal values of each pair of neurons in the whole network. The parameters are set as *k* = 1, *l* = 5, Δ*t* = 1 ms, and *m* = 1. The colors are the same as those in Fig. 3. (*G*) The similarity matrix of TE values over all neuron pairs across different stimuli conditions. For each pair of stimuli conditions, the similarity between the inferred TE values is measured by their correlation coefficient. The similarity matrix is symmetric with respect to its diagonal of ones, and the lower triangular part of the matrix is shown in the heatmap.

We then apply the same ROC analysis to all the four causality measures when analyzing experimental data under different visual stimuli conditions. As shown in the insets of Figs. 5*C-F*, the distribution of TDCC, TDMI, GC, and TE values are close to each other, which again verifies their mathematical relations given by Theorems 1-4. Under the assumption of mixed log-normal distribution (*SI Appendix*, Figs. S9A-D), we first infer the ground truth of the network structural connectivity, and then evaluate the reconstruction performance of TDCC, TDMI, GC, and TE by the ROC curves as shown in Figs. 5*C-F* with AUC values greater than 0.95. To investigate the consistency of reconstruction across different stimuli conditions, we compute the correlation coefficient of causal values for each pair of stimuli. As shown in Fig. 5*G*, our reconstruction achieves relatively high consistency across conditions with correlation coefficient higher than 0.75. With causal values, we also infer the binary adjacency matrix under each of four different stimuli conditions using the optimal inference threshold (see *Materials and Methods*) from the ROC analysis. Note that almost all of the causal values of inconsistently inferred connections fall into the overlapping region of the connected and unconnected distributions, *i*.*e*., the causal values from connected (unconnected) pairs but less (greater) than the threshold (green curve in *SI Appendix*, Figs. S9E-H) which are generally error prone. We point out that such error prone generally exists as we can see similar phenomena for HH networks (*SI Appendix*, Fig. S10).

## Discussion

In this work, inspired by the *pulse-output* nature of spiking neuronal networks, we have built an analytical framework upon the stochastic binary representation of neuronal interactions. It allows us, using only pairwise information between neurons, to (i) establish the mathematical relationships between four widely used measures (TDCC, TDMI, GC, and TE) of causal connectivity, and (ii) accurately reconstruct physical (structural) connectivity from causal connectivity in simulated HH neuronal networks. In addition, our framework provides the clarification and reduction of the major possible obstacles induced by confounders and unmeasured hidden nodes in conventional network reconstructions. Finally, we have used this analysis of pulse-output signals to reconstruct the structural connectivity of the real neuronal network in mouse brain from experimentally measured spike-train data and have achieved promising performances.

We emphasize two key features of pulse-output signals in eliminating of the curse of dimensionality that lead to effective network reconstruction via our framework: (i) the short timescale of auto-correlation, and (ii) the weakness of indirect causalities. On the one hand, with small time step Δ*t* commonly used in experiment, the short auto-correlation timescale protects the inferred causality from the corruption of the self-memory of time series. Therefore, the short auto-correlation length overcomes the curse of dimensionality in the estimation of probability density function by reducing the order of conditioned signal history (e.g., *k* = *l* = 1). On the other hand, because of the spike nature of the pulse-output signal, the causality value of direct connection is several orders of magnitude larger than that of indirect connection, which distinguishes direct connections from indirect ones. These two features make our framework a practical approach for pulse-coupled network reconstruction. In contrast, if these causality measures are directly applied to continuous-valued signals, *e*.*g*., voltage time series of a neuronal network, the mathematical relations derived in our theorems do not hold (*SI Appendix*, Fig. S11A) and the network reconstruction may also fail. For instance, TDCC and TDMI gives incorrect reconstruction of the structural connectivity due to the strong self-correlation of continuous-valued time series (*SI Appendix*, Fig. S11B).

In the main text, we have illustrated the effectiveness of the four causality measures by taking the examples of an excitatory HH neural network receiving uncorrelated external Poisson drive (see *Materials and Methods*). In fact, as discussed below, these methods apply much more broadly for pulse-output networks, including networks in different dynamical states, networks receiving correlated inputs, networks with both excitatory and inhibitory neurons, networks with different neuronal models.

First, oscillations and synchronizations are commonly observed in the biological brain network, as shown in *SI Appendix*, Fig. S12A. Due to the fake causality between neurons introduced by the strong synchronous state, conventional reconstruction frameworks fail to capture the true structural connectivity. However, with our framework, high inference accuracy (AUC *>* 0.88) can still be achieved (*SI Appendix*, Figs. S12B-C). Furthermore, by applying a desynchronized sampling method that only samples the pulse-output signals in asynchronous time intervals (*SI Appendix*, Fig. S12D), we can again perfectly reconstruct the network (AUC *>* 0.99, *SI Appendix*, Figs. S12E-F). The relation between percentages of desynchronized downsampling and AUC values is shown in *SI Appendix*, Fig. S13.

Second, external inputs to the network in the brain can often be correlated. In such a case, the synchronized states may be observed similarly as in previous cases (*SI Appendix*, Fig. S14A). Nevertheless, our framework can still achieve high inference accuracy (AUC *>* 0.99 with desynchronized downsampling methods, or AUC *>* 0.88 without downsampling, *SI Appendix*, Figs. S14 B-F) if the external inputs are moderately correlated, *e*.*g*., correlation coefficient less than 0.35 in our simulation case.

Third, in the case of HH networks with both excitatory and inhibitory neurons, we obtain similarly accurate reconstructions (AUC *>* 0.99 with desynchronized downsampling methods, or AUC *>* 0.71 without downsampling, *SI Appendix*, Fig. S15).

Last but not least, we also apply our framework of reconstruction to other types of neuronal networks, including leaky integrate-and-fire (LIF) networkAbbott (1999), Izhikevich networkIzhikevich (2003), FitzHugh-Nagumo networkFitzHugh (1961), and Morris-Lecar networkMorris and Lecar (1981) (see *Materials and Methods*). The results of all these networks can be seen in *SI Appendix*, Fig. S16. Our framework works well, with clear two-modal distributions of causality values (*SI Appendix*, Fig. S16B, S16E, S16H, S16K) and high reconstruction performance (AUC *>* 0.98, *SI Appendix*, Fig. S16C, S16F, S16I, S16L).

## Materials and Mechods

### Reconstruction performance evaluation

For binary inference of structural connectivity, analysis based on receiver operating characteristic (ROC) curves is adopted to evaluate the reconstruction performance in this work. The following two scenarios are considered.

#### With known structural connectivity

The conventional procedures for ROC curves analysis can be naturally applied towards data with true labels, i.e. structural connectivity in our binary reconstruction case. The area under the ROC curve (AUC) quantifies how well causality measures can distinguish structurally connected node pairs from unconnected node pairs. If AUC is close to 1, the distribution of the causality values of directly connected node pairs and that of unconnected ones are well distinguishable, i.e., the performance of binary reconstruction is good. If the AUC is close to 0.5, the distribution of those two kinds of causality values are virtually indistinguishable, meaning the performance of reconstruction is close to a random guess.

#### Without known structural connectivity

Conceptually, the causality measures for synaptically connected and unconnected pairs of neurons can be fitted by the log-normal distribution with different parameters, *i*.*e*., 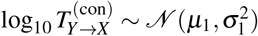 and 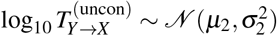 where the superscripts “con” and “uncon” indicates a direct synaptic connection and no synaptic connection from *Y* to *X*, respectively. So motivated, we fit the overall distribution with the summation of two log-normal distributions

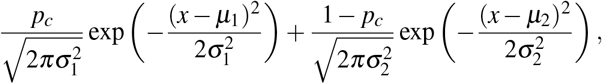

where *p*_*c*_ is the proportional coefficient between two Gaussian distributions 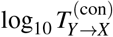 and 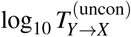. After that, the fitted distribution of two types of pairs are regarded as the true labels of structural connectivity. The similar procedures, as the previous case with known structural connectivity, are applied to obtain the ROC curve and the AUC value to evaluate the reconstruction performance.

### HH model

The dynamics of the *i*th neuron of an HH network is governed by Sun et al. (2009)

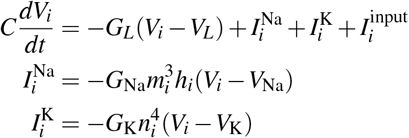

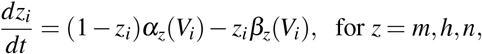

where *C* and *V*_*i*_ are the neuron’s membrane capacitance and membrane potential (voltage), respectively; *m*_*i*_, *h*_*i*_, and *n*_*i*_ are gating variables; *V*_Na_,*V*_K_, and *V*_L_ are the reversal potentials for the sodium, potassium, and leak currents, respectively; *G*_Na_, *G*_K_, and *G*_L_ are the corresponding maximum conductances; and *α*_*z*_ and *β*_*z*_ are the rate variables. The detailed dynamics of the gating variables *m, h, n* and the choice of parameters can be found in Ref. Dayan and Abbott (2001) and *SI Appendix, Supporting Information Text 3*. The input current 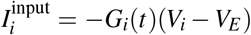 where *G*_*i*_(*t*) is the input conductance defined by *G*_*i*_(*t*) = *f* ∑_*l*_ *H*(*t* − *s*_*il*_) +∑ _*j*_ *A*_*i j*_*S* ∑_*l*_ *H*(*t* − *τ* _*jl*_) and *V*_*E*_ is the reversal potential of excitation. Here, *s*_*il*_ is the *l*th spike time of the external Poisson input with strength *f* and rate *ν*, **A** = (*A*_*i j*_) is the adjacency matrix with *A*_*i j*_ = 1 indicating a direct connection from neuron *j* to neuron *i* and *A*_*i j*_ = 0 indicating no connection there. *S* is the coupling strength, and *τ* _*jl*_ is the *l*th spike time of the *j*th neuron. The spike-induced conductance change *H*(*t*) is defined by 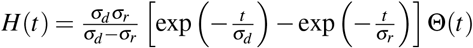, where *σ*_*d*_ and *σ*_*r*_ are the decay and rise time scale, respectively, and Θ (·) is the Heaviside function. When the voltage *V*_*j*_ reaches the firing threshold, *V*_th_, the *j*th neuron generates a spike at this time, say *τ* _*jl*_, and it will induce the *i*th neuron’s conductance change if *A*_*i j*_ = 1.

### LIF model

The dynamics of the *i*th neuron in a leaky integrate-and-fire (LIF) network is governed by Newhall et al. (2010); Gu et al. (2018)

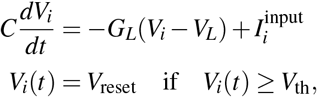

where *C* and *V*_*i*_ are the membrane capacitance and membrane potential. *V*_*L*_ and *G*_*L*_ are the reversal potential and conductance for leak currents. Compared with HH model, LIF model drops terms of nonlinear sodium and potassium current, and the input current is given by 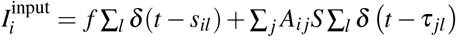, where *δ* (·) is the Dirac delta function, *s*_*il*_ is the *l*th spike time of the external Poisson input with strength *f*, and rate *ν, τ* _*jl*_ is the *l*th spike time of *j*th neuron with strength *S*. And **A** = (*A*_*i j*_) is the adjacency matrix defined the same as that in the HH model. When the voltage reaches the threshold *V*_th_, the *i*th neuron will emit a spike to all its connected post-synaptic neurons, and then reset to *V*_reset_ immediately. In numerical simulation, we use quantities: *C* = 1, *V*_reset_ = −65 mV, *V*_th_ = −40 mV, and the leakage conductance is set to be *G*_*L*_ = 0.05 ms^−1^ corresponding to the membrane time constant of 20 ms.

### Izhikevich model

The dynamics of the *i*th neuron in an Izhikevich network is governed by Izhikevich (2003)

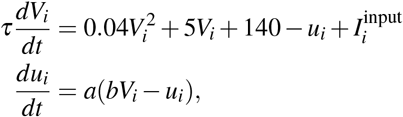

where *V*_*i*_ is the membrane potential and *u*_*i*_ is the recovery variable describing the force that drives *V*_*i*_ towards resting state. *τ* is the time constant of *V*_*i*_. *a* and *b* describe the time scale and sensitivity (with respect to *V*_*i*_) of *u*_*i*_. 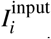 is the driving current containing the external Poisson input and the synaptic input from other neurons, defined by 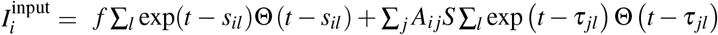, where Θ(·) is the Heaviside function and parameters are defined similarly as those in the LIF model. When the voltage reaches the threshold *V*_th_, the neuron emits a spike to all its post-synaptic neurons, and then reset *V*_*i*_ to *c*, and *u*_*i*_ to *u*_*i*_ + *d*. In numerical simulation, we use quantities: *τ* = 1 ms, *a* = 0.02 ms^−1^, *b* = 0.2 mV^−1^, *c* = −65 mV, *d* = 8, *V*_th_ = 30 mV.

### FitzHugh-Nagumo model

The dynamics of the *i*th neuron in a FitzHugh-Nagumo network is governed by FitzHugh (1961)

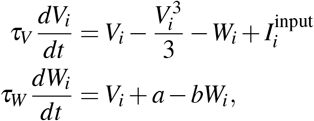

where *V*_*i*_ and *W*_*i*_ describe the membrane potential and recovery variable, respectively. *τ*_*V*_ and *τ*_*W*_ are their corresponding time constants. 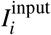 is the driving current containing the external Poisson input and the synaptic input from other neurons, defined by 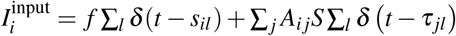, where parameters are defined similarly as those in the LIF model. When the voltage reaches the threshold *V*_th_, the neuron emits a spike to all its post-synaptic neurons.

In numerical simulation, we set quantities: *a* = 0.7, *b* = 0.8, *V*_th_ = 0, *τ*_*V*_ = 1 ms, and *τ*_*W*_ = 12.5 ms.

### Morris-Lecar model

The dynamics of the *i*th neuron in a Morris-Lecar network is governed by Morris and Lecar (1981)

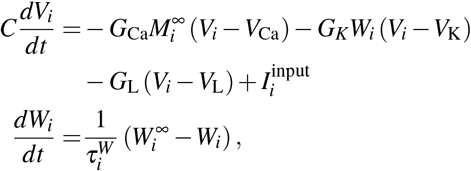

where *C* and *V*_*i*_ are the neuron’s membrane capacitance and membrane potential, respectively; *V*_Ca_,*V*_K_, and *V*_L_ are the reversal potentials for the calcium, potassium, and leak currents, respectively; *G*_Ca_, *G*_K_ and *G*_L_ are the corresponding maximum conductances; and *W*_*i*_ is the neuron’s recovery variable (normalized *K*^+^ conductance). 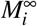 and 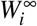 are the voltage-dependent equilibrium value of the normalized conductance of calcium and potassium, respectively, defined by

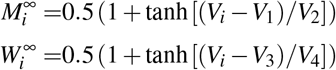

where *V*_1_, *V*_2_, *V*_3_, and *V*_4_ are the constant parameters. 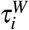 is a voltage-dependent time constant of *W*_*i*_, defined by

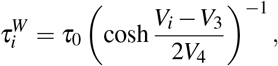

where *τ*_0_ is a temperature-dependent parameter, fixed as a constant in simulation. 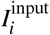 is the driving current containing the external Poisson input and the synaptic input from other neurons, defined by 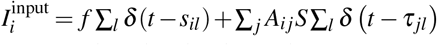, where parameters are defined similarly as those in LIF model. When the voltage reaches threshold *V*_th_, the neuron emits a spike to all its post-synaptic neurons. In numerical simulation, we set parameters as: *G*_Ca_ = 4 mS · cm^−2^, *V*_Ca_ = 120 mV, *G*_K_ = 8 mS · cm^−2^, *V*_K_ = −80 mV, *G*_L_ = 2 mS · cm^−2^, *V*_L_ = −60 mV, *C* = 20 mF · cm^−2^, *V*_1_ = −1.2 mV, *V*_2_ = 18 mV, *V*_3_ = 12 mV, *V*_4_ = 17.4 mV, *τ*_0_ = 15 ms, and *V*_th_ = 0 mV.

### Neurophysiological data

The public spike train data is from Allen brain observatory Allen Institute for Brain Science (2016), accessed via the Allen Software Development Kit (AllenSDK) Allen Institute for Brain Science (2018). Specifically, the data labeled with sessions-ID 715093703 was analyzed in this work. The 118-day-old male mouse passively received multiple visual stimuli from one of four categories, including drift gratings, static gratings, natural scenes and natural movies. The single neuronal activities, i.e. spike trains, were recorded from multiple brain areas, including APN, CA1, CA3, DG, LGd, LP, PO, VISam, VISl, VISp, VISpm, and VISrl, using 6 Neuropixel probes. For each category of stimulus, the recording lasts for more than 20 minutes. 884 sorted spike trains were recorded and 156 of those were used for causality analysis with signal-to-noise ratio greater than 4 and mean firing rate greater than 0.08 Hz.

## Data Availability

Implementation of causality estimation, network simulation, and processing scripts for Allen Brain Observatory data are all available through GitHub (https://github.com/neoneuron/causal4).

## Acknowledgements

This work was supported by Science and Technology Innovation 2030 - Brain Science and Brain-Inspired Intelligence Project with Grant No. 2021ZD0200204 and the Lingang Laboratory Grant No. LG-QS-202202-01 (S.L., D.Z.,); National Natural Science Foundation of China Grant 12271361 (S.L.); National Natural Science Foundation of China with Grant No. 12071287, 12225109 (D.Z.), Shanghai Municipal Science and Technology Major Project 2021SHZDZX0102 and the Student Innovation Center at Shanghai Jiao Tong University (Z.K.T, K.C., S.L., D.Z.)

