## Supplementary Information for "Quantitative relations among causality measures with applications to pulse-output nonlinear network reconstruction"

#### 1 Mathematical derivation of relations among four causality measures

We first derive the mathematical relations among time-delayed correlation coefficient (TDCC), time-delayed mutual information (TDMI), Granger causality (GC), and transfer entropy (TE) for networks with pulse signals as measured output. Consider a pair of nodes in a network, say nodes $X$ and $Y$ with pulse-output signals $w_x(t) = \sum_l \delta(t - \tau_{xl})$ and $w_y(t) = \sum_l \delta(t - \tau_{yl})$, where $\delta(\cdot)$ is the Dirac delta function. Under a sampling resolution $\Delta t$, the pulse-output signals are measured as binary time series $\{x_n\}$ and $\{y_n\}$, where $x_n = 1$ ($y_n = 1$) if there is a pulse signal of $X$ ($Y$) in the time window $[t_n, t_{n+1})$, and $x_n = 0$ ($y_n = 0$) otherwise, *i.e.*,

$$x_n = \int_{t_n}^{t_{n+1}} w_x(t)dt \quad \text{and} \quad y_n = \int_{t_n}^{t_{n+1}} w_y(t)dt,$$

and $t_n = n\Delta t$. Note that the value of $\Delta t$ is chosen to be small to make sure that there is at most one pulse signal in one time window. The responses $x_n$ and $y_n$ are viewed as stochastic processes, as they are when the network is driven by stationary stochastic inputs. Accordingly, below we will describe the neuronal responses probabilistically.

For the ease of discussion, we define the following notations:

$$r_x = \frac{1}{T}\int_0^T w_x(t)dt \quad \text{and} \quad r_y = \frac{1}{T}\int_0^T w_y(t)dt$$

are the mean pulse rates of $X$ and $Y$, respectively;

$$p_x = p(x_n = 1) \quad \text{and} \quad p_y = p(y_n = 1)$$

are the probability of $x_n$ and $y_n$ being 1, respectively. Then we have

$$p_x = r_x\Delta t = O(\Delta t), \;\; p_y = r_y\Delta t = O(\Delta t) \tag{1}$$

and

$$\sigma_x^2 = p_x - p_x^2 = O(\Delta t), \;\; \sigma_y^2 = p_y - p_y^2 = O(\Delta t),$$

where the symbol "$O$" stands for the order, $\sigma_x$ and $\sigma_y$ are the standard deviation of $\{x_n\}$ and $\{y_n\}$, respectively. Also, we define $\Delta p(x_n, y_{n-m})$ measuring the dependence between $x_n$ being $\xi$ and $y_{n-m}$ being $\eta$ by

$$\Delta p(x_n = \xi, y_{n-m} = \eta) = \frac{p(x_n = \xi, y_{n-m} = \eta)}{p(x_n = \xi)p(y_{n-m} = \eta)} - 1. \tag{2}$$

Specially, we denote the dependence between $x_n$ and $y_{n-m}$ being 1 by

$$\Delta p_m = \Delta p(x_n = 1, y_{n-m} = 1) = \frac{p(x_n = 1, y_{n-m} = 1)}{p(x_n = 1)p(y_{n-m} = 1)} - 1. \tag{3}$$

#### 1.1 Definition of TDCC, TDMI, GC, and TE

Without loss of generality, we consider the causal interaction from $Y$ to $X$ with binary time series $\{x_n\}$ and $\{y_n\}$. TDCC from $Y$ to $X$ is defined by

$$C(X,Y;m) = \frac{\mathrm{cov}(x_n, y_{n-m})}{\sigma_x \sigma_y}, \tag{4}$$

where $m > 0$ is the time delay.

TDMI from $Y$ to $X$ is defined by

$$I(X,Y;m) = \sum_{x_n, y_{n-m}} p(x_n, y_{n-m}) \log \frac{p(x_n, y_{n-m})}{p(x_n)p(y_{n-m})},$$

where $p(x_n, y_{n-m})$ is the joint probability distribution of $x_n$ and $y_{n-m}$, $p(x_n)$ and $p(y_{n-m})$ are the corresponding marginal probability distributions.

GC is established based on linear regression. The auto-regression for $X$ is represented by

$$x_{n+1} = a_0 + \sum_{i=1}^{k} a_i x_{n+1-i} + \varepsilon_{n+1},$$

where $\{a_i\}$ are the auto-regression coefficients and $\varepsilon_{n+1}$ is the residual. By including the historical information of $Y$ with message length $l$ and time delay $m$, the joint regression for $X$ is represented by

$$x_{n+1} = \tilde{a}_0 + \sum_{i=1}^{k} \tilde{a}_i x_{n+1-i} + \sum_{j=1}^{l} \tilde{b}_j y_{n+2-m-j} + \eta_{n+1},$$

where $\{\tilde{a}_i\}$ and $\{\tilde{b}_j\}$ are the joint regression coefficients, and $\eta_{n+1}$ is the corresponding residual. The GC value from $Y$ to $X$ is defined by

$$G_{Y\to X}(k,l;m) = \log \frac{\mathrm{Var}(\varepsilon_{n+1})}{\mathrm{Var}(\eta_{n+1})}.$$

By introducing the time-delay parameter $m$, the GC analysis defined above generalizes the conventional GC analysis, as the latter corresponds to the special case of $m = 1$.

The TE value from $Y$ to $X$ is defined by

$$T_{Y\to X}(k,l;m) = \sum_{x_{n+1}, x_n^{(k)}, y_{n+1-m}^{(l)}} p(x_{n+1}, x_n^{(k)}, y_{n+1-m}^{(l)}) \log \frac{p(x_{n+1}|x_n^{(k)}, y_{n+1-m}^{(l)})}{p(x_{n+1}|x_n^{(k)})}, \tag{5}$$

where the shorthand notation $x_n^{(k)} = (x_n, x_{n-1}, ..., x_{n-k+1})$ and $y_{n+1-m}^{(l)} = (y_{n+1-m}, y_{n-m}, ..., y_{n+2-m-l})$, $k$, $l$ indicate the length (order) of historical information of $X$ and $Y$, respectively. Similar to GC, the time-delay parameter $m$ is introduced that generalizes the conventional TE, the latter of which corresponds to the case of $m = 1$.

#### 1.2 Mathematical relation between TDMI and TDCC

From the definition of TDCC in Eq. 4, for binary value time series $\{x_n\}$ and $\{y_{n-m}\}$, we have

$$\begin{aligned} C(X,Y;m) &= \frac{\operatorname{cov}(x_n, y_{n-m})}{\sigma_x \sigma_y} \\ &= \frac{E[(x_n - E[x_n])(y_{n-m} - E[y_{n-m}])]}{\sqrt{p_x(1-p_x)}\sqrt{p_y(1-p_y)}} \\ &= \frac{E[(x_n - p_x)(y_{n-m} - p_y)]}{\sqrt{p_x(1-p_x)}\sqrt{p_y(1-p_y)}} \\ &= \frac{p(x_n = 1, y_{n-m} = 1) - p_x p_y}{\sqrt{(p_x - p_x^2)(p_y - p_y^2)}}. \end{aligned} \tag{6}$$

The relation between TDMI and TDCC can be derived by Taylor expanding TDMI with respect to $\Delta p(x_n, y_{n-m})$, defined in Eq. 2, as follows:

$$\begin{aligned} I(X,Y;m) &= \sum_{x_n, y_{n-m}} p(x_n)p(y_{n-m}) \left[1 + \left(\frac{p(x_n, y_{n-m})}{p(x_n)p(y_{n-m})} - 1\right)\right] \log\left[1 + \left(\frac{p(x_n, y_{n-m})}{p(x_n)p(y_{n-m})} - 1\right)\right] \\ &= \sum_{\xi, \eta \in \{0,1\}} p(x_n = \xi)p(y_{n-m} = \eta)\,[1 + \Delta p(x_n = \xi, y_{n-m} = \eta)] \log[1 + \Delta p(x_n = \xi, y_{n-m} = \eta)] \\ &= \sum_{\xi, \eta \in \{0,1\}} p(x_n = \xi)p(y_{n-m} = \eta) \Big[\Delta p(x_n = \xi, y_{n-m} = \eta) + \frac{1}{2}\Delta p^2(x_n = \xi, y_{n-m} = \eta) \\ &\qquad + O\left(\Delta p^3(x_n = \xi, y_{n-m} = \eta)\right)\Big] \end{aligned} \tag{7}$$

Due to the simplicity of binary value series, the summation in Eq. 7 contains only four terms. We list the expression of $\Delta p(x_n, y_{n-m})$ for all possible $\xi$ and $\eta$ values in Table 1 in terms of $\Delta p_m$, defined by Eq. 3.

| $\Delta p(x_n, y_{n-m})$ | $x_n = 0$ | $x_n = 1$ |
|---|---|---|
| $y_{n-m} = 0$ | $\frac{p_x p_y \Delta p_m}{(1-p_x)(1-p_y)}$ | $-\frac{p_x p_y \Delta p_m}{p_x(1-p_y)}$ |
| $y_{n-m} = 1$ | $-\frac{p_x p_y \Delta p_m}{(1-p_x)p_y}$ | $\Delta p_m$ |

Table 1: Expressions of $\Delta p(x_n, y_{n-m})$ in terms of $\Delta p_m$.

Then, we substitute all four terms of $\Delta p(x_n, y_{n-m})$ in Eq. 7 to obtain

$$\begin{aligned} I(X,Y;m) &= \frac{[p(x_n = 1, y_{n-m} = 1) - p_x p_y]^2}{2(p_x - p_x^2)(p_y - p_y^2)} + O(\Delta t^2 \Delta p_m^3) \\ &= \frac{C^2(X,Y;m)}{2} + O(\Delta t^2 \Delta p_m^3). \end{aligned} \tag{8}$$

#### 1.3 Mathematical relation between GC and TDCC

From the definition, GC can be represented by the covariances of the signals as Barnett et al. (2009)

$$G_{Y\to X}(k,l;m) = \log\frac{\Gamma(x_{n+1}|x_n^{(k)})}{\Gamma(x_{n+1}|x_n^{(k)}\oplus y_{n+1-m}^{(l)})}, \tag{9}$$

where $\Gamma(\mathbf{x}|\mathbf{y}) = \mathrm{cov}(\mathbf{x}) - \mathrm{cov}(\mathbf{x},\mathbf{y})\mathrm{cov}(\mathbf{y})^{-1}\mathrm{cov}(\mathbf{x},\mathbf{y})^T$ for random vectors $\mathbf{x}$ and $\mathbf{y}$, $\mathrm{cov}(\mathbf{x})$ and $\mathrm{cov}(\mathbf{y})$ denote the covariance matrix of $\mathbf{x}$ and $\mathbf{y}$, respectively, and $\mathrm{cov}(\mathbf{x},\mathbf{y})$ denotes the cross-covariance matrix between $\mathbf{x}$ and $\mathbf{y}$. The symbol $T$ is the transpose operator and $\oplus$ denotes the concatenation of vectors.

To derive the relation between GC and TDCC, we first consider the auto-correlation function (ACF) of the binary time series $\{x_n\}$ and $\{y_n\}$. The ACF of $\{x_n\}$ is defined as

$$\mathrm{ACF}(X;m) = \frac{\mathrm{cov}(x_n,x_{n-m})}{\sigma_x^2},$$

where $m$ is the non-zero time delay. Let $g_x(t)$ be the probability density function that node $X$ will generate a pulse at time $t$ given that it produced a pulse at time $t=0$. Then we have

$$p(x_n = 1|x_{n-m} = 1) = g_x(m\Delta t)\Delta t + O(\Delta t^2).$$

In general, the function $g_x(t)$ is continuous and bounded, thus we have $p(x_n = 1|x_{n-m} = 1) = O(\Delta t)$. Together with Eq. 1, we can obtain

$$\mathrm{ACF}(X;m) = \frac{p(x_n = 1|x_{n-m} = 1) - p_x}{1-p_x} = O(\Delta t). \tag{10}$$

Similarly, we have

$$\mathrm{ACF}(Y;m) = O(\Delta t).$$

Based on this, we derive the relation between GC and TDCC as follows: from Eq. 10, we can obtain

$$\mathrm{cov}(x_n^{(k)}) = \sigma_x^2(\mathbf{I}+\hat{\mathbf{A}}),$$

$$\mathrm{cov}(x_n^{(k)})^{-1} = \frac{1}{\sigma_x^2}(\mathbf{I}-\hat{\mathbf{A}}) + O(\Delta t\mathbf{1}_{k\times k}),$$

where $\hat{\mathbf{A}} = (\hat{a}_{ij})$, $\hat{a}_{ij} = O(\Delta t)$. Note that $\sigma_x^2 = O(\Delta t)$, thus the order with respect to $\Delta t$ in first term of the covariance matrix is up to $O(1)$. Besides, $\mathbf{I}$ is the identity matrix, and $\mathbf{1}_{k\times k}$ is the all-one matrix. Thus,

$$\Gamma(x_{n+1}|x_n^{(k)}) = \sigma_x^2 - \frac{1}{\sigma_x^2}\mathrm{cov}(x_{n+1},x_n^{(k)})(\mathbf{I}-\hat{\mathbf{A}})\mathrm{cov}(x_{n+1},x_n^{(k)})^T + O(\Delta t^5). \tag{11}$$

In the same way, we have

$$\mathrm{cov}(x_n^{(k)}\oplus y_{n+1-m}^{(l)}) = \begin{pmatrix} \sigma_x^2(\mathbf{I}+\hat{\mathbf{A}}) & \sigma_x\sigma_y\hat{\mathbf{C}} \\ \sigma_x\sigma_y\hat{\mathbf{C}}^T & \sigma_y^2(\mathbf{I}+\hat{\mathbf{B}}) \end{pmatrix},$$

$$\mathrm{cov}(x_n^{(k)} \oplus y_{n+1-m}^{(l)})^{-1} = \begin{pmatrix} (\mathbf{I}-\hat{\mathbf{A}})/\sigma_x^2 & -\hat{\mathbf{C}}/\sigma_x\sigma_y \\ -\hat{\mathbf{C}}^T/\sigma_x\sigma_y & (\mathbf{I}-\hat{\mathbf{B}})/\sigma_y^2 \end{pmatrix} + O\left(\Delta t \mathbf{1}_{(k+l)\times(k+l)}\right),$$

where $\hat{\mathbf{B}} = (\hat{b}_{ij})$, $\hat{b}_{ij} = O(\Delta t)$, $\hat{\mathbf{C}} = (\hat{c}_{ij})$, $\hat{c}_{ij} = O(\Delta t \Delta p_m)$. Similarly since $\sigma_x^2$ and $\sigma_y^2$ is $O(\Delta t)$, the first term of the inverse of covariance matrix is $O(1)$ with respect to $\Delta t$. Thus,

$$\begin{aligned}\Gamma(x_{n+1}|x_n^{(k)} \oplus y_{n+1-m}^{(l)}) =& \sigma_x^2 - \frac{1}{\sigma_x^2}\mathrm{cov}(x_{n+1},x_n^{(k)})(\mathbf{I}-\hat{\mathbf{A}})\,\mathrm{cov}(x_{n+1},x_n^{(k)})^T \\ & - \frac{1}{\sigma_y^2}\mathrm{cov}(x_{n+1},y_{n+1-m}^{(l)})(\mathbf{I}-\hat{\mathbf{B}})\,\mathrm{cov}(x_{n+1},y_{n+1-m}^{(l)})^T \\ & + \frac{2}{\sigma_x\sigma_y}\mathrm{cov}(x_{n+1},x_n^{(k)})\hat{\mathbf{C}}\,\mathrm{cov}(x_{n+1},y_{n+1-m}^{(l)})^T + O(\Delta t^5).\end{aligned} \tag{12}$$

Substituting Eqs. 11 and 12 into Eq. 9 and Taylor expanding Eq. 9 with respect to $\Delta t$, we can obtain

$$\begin{aligned}G_{Y\to X}(k,l;m) =& \frac{\mathrm{cov}(x_{n+1},y_{n+1-m}^{(l)})\,\mathrm{cov}(x_{n+1},y_{n+1-m}^{(l)})^T}{\sigma_x^2\sigma_y^2} \\ & \underbrace{-\frac{1}{\sigma_x^2\sigma_y^2}\left[\mathrm{cov}(x_{n+1},y_{n+1-m}^{(l)})\hat{\mathbf{B}}\,\mathrm{cov}(x_{n+1},y_{n+1-m}^{(l)})^T + \frac{2\sigma_y}{\sigma_x}\mathrm{cov}(x_{n+1},x_n^{(k)})\hat{\mathbf{C}}\,\mathrm{cov}(x_{n+1},y_{n+1-m}^{(l)})^T\right]}_{O(\Delta t^3 \Delta p_m^2)} \\ & + O(\Delta t^4 \Delta p_m^4)\end{aligned} \tag{13}$$

Note that the first term in Eq. 13 is the cross correlation between $x_{n+1}$ and $y_{n+1-m}^{(1)}$, thus, by dropping the higher order term $O(\Delta t^4)$, we have

$$G_{Y\to X}(k,l;m) = \sum_{i=m}^{m+l-1} C^2(X,Y;i) + O(\Delta t^3 \Delta p_m^2). \tag{14}$$

#### 1.4 Mathematical relation between TE and TDMI

To rigorously establish the relation between TE and TDMI, we require that $\left\|x_{n+1}^{(k+1)}\right\|_0 \le 1$ and $\left\|y_{n+1-m}^{(l)}\right\|_0 \le 1$ in the definition of TE given in Eq. 5, where $\left\|\cdot\right\|_0$ denotes the $l_0$ norm of a vector, *i.e.*, the number of nonzero elements in a vector. This assumption indicates that the length of historical information used in the TE framework is shorter than the "refractory period", *i.e.*, the minimal time interval between two consecutive pulse-output signals.

For simplify, we use $x^-$and $y^-$ to denote $x_n^{(k)} = (x_n, x_{n-1}, \cdots, x_{n-k+1})$ and $y_{n+1-m}^{(l)} = (y_{n+1-m}, y_{n-m}, \cdots, y_{n+2-m-l})$, respectively. From the definition of TE, we have

$$\begin{aligned}T_{Y\to X}(k,l;m) &= \sum_{x_{n+1},x^-,y^-} p(x_{n+1},x^-,y^-)\log\frac{p(x_{n+1}|x^-,y^-)}{p(x_{n+1}|x^-)} \\ &= \sum_{x_{n+1},x^-,y^-} p(x_{n+1},x^-,y^-)\left[\log\frac{p(x_{n+1}|y^-)}{p(x_{n+1})} + \log\frac{p(x_{n+1}|x^-,y^-)}{p(x_{n+1}|y^-)}\frac{p(x_{n+1})}{p(x_{n+1}|x^-)}\right].\end{aligned}$$

Because
$$\begin{aligned}\sum_{x_{n+1},y^-} p(x_{n+1},y^-)\log\frac{p(x_{n+1}|y^-)}{p(x_{n+1})} &= \sum_{x_{n+1},y^-} p(x_{n+1},y^-)\log\frac{p(y^-|x_{n+1})}{p(y^-)} \\ &= \sum_{x_{n+1},y^-} p(x_{n+1},y^-)\left[\log\frac{\prod_j p(y_j|x_{n+1})}{\prod_j p(y_j)} + \log\frac{p(y^-|x_{n+1})}{\prod_j p(y_j|x_{n+1})}\frac{\prod_j p(y_j)}{p(y^-)}\right] \\ &= \sum_{i=m}^{m+l-1} I(X,Y;i) + \sum_{x_{n+1},y^-} p(x_{n+1},y^-)\log\frac{p(y^-|x_{n+1})}{\prod_j p(y_j|x_{n+1})}\frac{\prod_j p(y_j)}{p(y^-)},\end{aligned}$$

where $\prod_j$ represents $\prod_{j=n+2-m-l}^{n+1-m}$, we have

$$T_{Y\to X}(k,l;m) = \sum_{i=m}^{m+l-1} I(X,Y;i) + \mathscr{A} + \mathscr{B}, \tag{15}$$

where
$$\mathscr{A} = \sum_{x_{n+1},y^-} p(x_{n+1},y^-)\log\frac{p(y^-|x_{n+1})}{\prod_j p(y_j|x_{n+1})}\frac{\prod_j p(y_j)}{p(y^-)}$$
and
$$\mathscr{B} = \sum_{x_{n+1},x^-,y^-} p(x_{n+1},x^-,y^-)\log\frac{p(x_{n+1}|x^-,y^-)}{p(x_{n+1}|y^-)}\frac{p(x_{n+1})}{p(x_{n+1}|x^-)}.$$

Under the assumption that $\left\|x_{n+1}^{(k+1)}\right\|_0 \le 1$ and $\left\|y_{n+1-m}^{(l)}\right\|_0 \le 1$, the number of nonzero components is at most one in $x_{n+1}^{(k+1)}$ and $y_{n+1-m}^{(l)}$. We use $1_{x_s}$ to denote the event that only the state $x_s$ is one in $x^-$, where $n-k+1 \le s \le n$, and use $0_x^-$ to denote the event that all the components in $x^-$ are zero. Similarly, we use $1_{y_t}$ to denote the event that only the state $y_t$ is one in $y^-$, where $n+2-m-l \le t \le n+1-m$, and use $0_y^-$ to denote the event that all the components in $y^-$ are zero. Then we can derive the leading order of each term in $\mathscr{A}$ and $\mathscr{B}$ by Taylor expanding them with respect to $\Delta t$ and $\Delta p_m$.

In $\mathscr{A}$, we define the dependence between $x_{n+1}$ and $y^-$, similarly as in Eqs. 2 and 3, by
$$\Delta p(x_{n+1},y^-) = \frac{p(x_{n+1},y^-)}{p(x_{n+1})p(y^-)} - 1.$$
And more specifically, we define
$$\Delta p_{n+1-t} = \Delta p(x_{n+1}=1, y^- = 1_{y_t}) = \frac{p(x_{n+1}=1, y^- = 1_{y_t})}{p(x_{n+1}=1)p(y^- = 1_{y_t})} - 1, \tag{16}$$
where $n+2-m-l \le t \le n+1-m$. Then, we can construct the table of $p(x_{n+1},y^-)$ in terms of $\Delta p_{n+1-t}$, shown in Table 2.

For the terms in $\mathscr{A}$ of which $x_{n+1}=1$ and $y^- = 1_{y_t}$,
$$p(x_{n+1},y^-)\log\frac{p(y^-|x_{n+1})}{\prod_j p(y_j|x_{n+1})}\frac{\prod_j p(y_j)}{p(y^-)}\Bigg|_{x_{n+1}=1,y^-=1_{y_t}} \tag{17}$$
where $\prod_j$ represents $\prod_{j=n+2-m-l}^{n+1-m}$, we have

$$p(x_{n+1}=1, y_t=1) = p(x_{n+1}=1, y^- = 1_{y_t}) = p_x p_y(1+\Delta p_{n+1-t}), \tag{18}$$

| $p(x_{n+1}, y^-)$ | $x_{n+1} = 0$ | $x_{n+1} = 1$ |
|---|---|---|
| $y^- = 0_y^-$ | $1 - p_x - lp_y + p_x p_y \left( l + \sum_{t=n+2-m-l}^{n-m+1} \Delta p_{n+1-t} \right)$ | $p_x - p_x p_y \left( l + \sum_{t=n+2-m-l}^{n-m+1} \Delta p_{n+1-t} \right)$ |
| $y^- = 1_{y_t}$ | $p_y - p_x p_y \left(1 + \Delta p_{n+1-t}\right)$ | $p_x p_y \Delta p_{n+1-t}$ |

Table 2: Expressions of $p(x_{n+1}, y^-)$ in terms of $\Delta p_{n+1-t}$, where $n+2-m-l \leq t \leq n+1-m$.

$$p(y_j = 0 | x_{n+1} = 1) = \frac{p(x_{n+1} = 1, y_j = 0)}{p(x_{n+1} = 1)} = \frac{p_x - p(x_{n+1} = 1, y_j = 1)}{p_x} = 1 - p_y(1 + \Delta p_{n+1-j}). \tag{19}$$

Substituting Eqs. 18-19 and corresponding entries in Table 2 into Eq. 17 yields

$$\begin{aligned}
& p(x_{n+1}, y^-) \log \frac{p(y^- | x_{n+1})}{\prod_j p(y_j | x_{n+1})} \frac{\prod_j p(y_j)}{p(y^-)} \Bigg|_{x_{n+1}=1, y^-=1_{y_t}} \\
&= p_x p_y (1 + \Delta p_{n+1-t}) \log \frac{(1-p_y)^{l-1}}{\prod_{j \neq t} (1 - p_y - p_y \Delta p_{n+1-j})} \\
&= p_x p_y (1 + \Delta p_{n+1-t}) \sum_{j \neq t} \log \frac{1 - p_y}{1 - p_y - p_y \Delta p_{n+1-j}} \\
&= p_x p_y^2 \sum_{j \neq t} \Delta p_{n+1-j} \underbrace{+ p_x p_y^2 \Delta p_{n+1-t} \sum_{j \neq t} \Delta p_{n+1-j}}_{O(\Delta t^3 \Delta p_m^2)} + O(\Delta t^4 \Delta p_m^2) \\
&= p_x p_y^2 \sum_{j \neq t} \Delta p_{n+1-j} + O(\Delta t^3 \Delta p_m^2).
\end{aligned} \tag{20}$$

Note that we take $\Delta p_{n+1-j} = O(\Delta p_m)$ in the above derivation. Other terms in $\mathscr{A}$ can be obtained similarly as follows:

$$\begin{aligned}
& p(x_{n+1}, y^-) \log \frac{p(y^- | x_{n+1})}{\prod_j p(y_j | x_{n+1})} \frac{\prod_j p(y_j)}{p(y^-)} \Bigg|_{x_{n+1}=1, y^-=0_y^-} \\
&= \left( p_x - p_x p_y (l + \sum_t \Delta p_{n+1-t}) \right) \log \frac{1 - p_y (l + \sum_t \Delta p_{n+1-t})}{\prod_t (1 - p_y - p_y \Delta p_{n+1-t})} \frac{(1-p_y)^l}{1 - lp_y} \\
&= (1-l) p_x p_y^2 \sum_t \Delta p_{n+1-t} \underbrace{+ \frac{1}{2} p_x p_y^2 \left[ \sum_j \Delta p_{n+1-j}^2 - \left( \sum_j \Delta p_{n+1-j} \right)^2 \right]}_{O(\Delta t^3 \Delta p_m^2)} + O(\Delta t^4),
\end{aligned} \tag{21}$$

$$
\begin{aligned}
&p(x_{n+1},y^-)\log\frac{p(y^-|x_{n+1})}{\prod_j p(y_j|x_{n+1})}\frac{\prod_j p(y_j)}{p(y^-)}\bigg|_{x_{n+1}=0,y^-=1_{y_t}} \\
&=(p_y-p_xp_y(1+\Delta p_{n+1-t}))\log\frac{(1-p_y)^{l-1}(1-p_x)^{l-1}}{\prod_{j\neq t}\big(1-p_x-p_y+p_xp_y(1+\Delta p_{n+1-j})\big)} \\
&=-p_xp_y^2\sum_{j\neq t}\Delta p_{n+1-j}+O(\Delta t^4),
\end{aligned}
\tag{22}
$$

$$
\begin{aligned}
&p(x_{n+1},y^-)\log\frac{p(y^-|x_{n+1})}{\prod_j p(y_j|x_{n+1})}\frac{\prod_j p(y_j)}{p(y^-)}\bigg|_{x_{n+1}=0,y^-=0_y^-} \\
&=\Big(1-p_x-lp_y+p_xp_y(l+\sum_t\Delta p_{n+1-t})\Big)\log\frac{1-p_x-lp_y+p_xp_y(l+\sum_t\Delta p_{n+1-t})}{\prod_t(1-p_x-p_y+p_xp_y(1+\Delta p_{n+1-t}))}\frac{(1-p_y)^l(1-p_x)^l}{(1-lp_y)(1-p_x)} \\
&=(l-1)p_xp_y^2\sum_t\Delta p_{n+1-t}+O(\Delta t^4).
\end{aligned}
\tag{23}
$$

Therefore, combining Eqs. 20-23, we obtain $\mathscr{A}=O(\Delta t^3\Delta p_m^2)$.

For

$$
\mathscr{B}=\sum_{x_{n+1},x^-}p(x_{n+1},x^-)\log\frac{p(x_{n+1})}{p(x_{n+1}|x^-)}+\sum_{x_{n+1},x^-,y^-}p(x_{n+1},x^-,y^-)\log\frac{p(x_{n+1}|x^-,y^-)}{p(x_{n+1}|y^-)},
\tag{24}
$$

the first term is the negative mutual information between $x_{n+1}$ and $x^-$. With the $\left\|x_n^{(k+1)}\right\|_0\leq 1$ assumption in **Theorem 3**, we write down the joint probability distribution in Table 3,

| $p(x_{n+1},x^-)$ | $x_{n+1}=0$ | $x_{n+1}=1$ |
|---|---|---|
| $x^-=0_x^-$ | $1-(1+k)p_x$ | $p_x$ |
| $x^-=1_{x_s}$ | $p_x$ | 0 |

Table 3: Expressions of $p(x_{n+1},x^-)$ in terms of $p_x$, where $n-k+1\leq s\leq n$.

And we can estimate the order of the first term by

$$
\begin{aligned}
&\sum_{x_{n+1},x^-}p(x_{n+1},x^-)\log\frac{p(x_{n+1})}{p(x_{n+1}|x^-)} \\
&=(1-(k+1)p_x)\log\frac{(1-p_x)(1-kp_x)}{1-(k+1)p_x}+p_x\log(1-kp_x)+\sum_s p_x\log(1-p_x) \\
&=(1-(k+1)p_x)\log\left(1+\frac{kp_x^2}{1-(k+1)p_x}\right)+p_x\log(1-kp_x)+kp_x\log(1-p_x) \\
&=-kp_x^2-\frac{k(k+1)}{2}p_x^3+O(\Delta t^4).
\end{aligned}
\tag{25}
$$

For the second term in Eq. 24, we consider the joint probability distribution $p(x_{n+1},x^-,y^-)$, and we define the dependence $\Delta p(x_{n+1},x^-,y^-)$ by

$$\Delta p(x_{n+1}, x^-, y^-) = \frac{p(x_{n+1}, x^-, y^-)}{p(x_{n+1})p(x^-, y^-)} - 1. \tag{26}$$

More specifically,

$$\begin{aligned}
\Delta p_{n+1-t} = \Delta p(x_{n+1}=1, x^-=0^-_x, y^-=1_{y_t}) &= \frac{p(x_{n+1}=1, x^-=0^-_x, y^-=1_{y_t})}{p(x_{n+1}=1, x^-=0^-_x)p(y^-=1_{y_t})} - 1 \\
&= \frac{p(x_{n+1}=1, x^-=0^-_x, y^-=1_{y_t})}{p_x p_y} - 1, \\
\Delta p_{s-t} = \Delta p(x_{n+1}=0, x^-=1_{x_s}, y^-=1_{y_t}) &= \frac{p(x_{n+1}=0, x^-=1_{x_s}, y^-=1_{y_t})}{p(x_{n+1}=0, x^-=1_{x_s})p(y^-=1_{y_t})} - 1 \\
&= \frac{p(x_{n+1}=0, x^-=1_{x_s}, y^-=1_{y_t})}{p_x p_y} - 1.
\end{aligned}$$

Then, we deduce the joint probability distribution $p(x_{n+1}, x^-, y^-)$ in terms of $\Delta p_{n+1-t}$ and $\Delta p_{s-t}$ as shown in Tables 4-5.

| $p(x_{n+1}, x^-=0^-_x, y^-)$ | $x_{n+1}=0$ | $x_{n+1}=1$ |
|---|---|---|
| $y^-=0^-_y$ | $1-(1+k)p_x - l p_y + p_x p_y \left((1+k)l + \sum_t \Delta p_{n+1-t} + \sum_{s,t} p_{s-t}\right)$ | $p_x - p_x p_y \left(l + \sum_t \Delta p_{n+1-t}\right)$ |
| $y^-=1_{y_t}$ | $p_y - p_x p_y \left(1 + \Delta p_{n+1-t} + k + \sum_s p_{s-t}\right)$ | $p_x p_y (1+\Delta p_{n+1-t})$ |

Table 4: Expressions of $p(x_{n+1}, x^-=0^-_x, y^-)$ in terms of $\Delta p_{s-t}$ and $\Delta p_{n+1-t}$, where $n-k+1 \le s \le n$ and $n+2-m-l \le t \le n+1-m$.

| $p(x_{n+1}, x^-=1_{x_s}, y^-)$ | $x_{n+1}=0$ | $x_{n+1}=1$ |
|---|---|---|
| $y^-=0^-_y$ | $p_x - p_x p_y \left(l + \sum_t \Delta p_{s-t}\right)$ | 0 |
| $y^-=1_{y_t}$ | $p_x p_y (1+\Delta p_{s-t})$ | 0 |

Table 5: Expressions of $p(x_{n+1}, x^-=1_{x_s}, y^-)$ in terms of $\Delta p_{s-t}$ and $\Delta p_{n+1-t}$, where $n-k+1 \le s \le n$ and $n+2-m-l \le t \le n+1-m$.

Finally, we write down all different types of terms in $\mathscr{B}$, with the help of tables above,

$$p(x_{n+1},x^-,y^-)\log\frac{p(x_{n+1}|x^-,y^-)}{p(x_{n+1}|y^-)}\Big|_{x_{n+1}=1,x^-=0_x^-,y^-=1_{y_t}}$$
$$=-p_xp_y(1+\Delta p_{n+1-t})\log\left(1-p_x(k+\sum_s\Delta p_{s-t})\right) \tag{27}$$
$$=p_x^2p_y(k+k\Delta p_{n+1-t}+\sum_s\Delta p_{s-t})\underbrace{+p_x^2p_y\Delta p_{n+1-t}\left(\sum_s\Delta p_{s-t}\right)}_{O(\Delta t^3\Delta p_m^2)}+O(\Delta t^4),$$

$$p(x_{n+1},x^-,y^-)\log\frac{p(x_{n+1}|x^-,y^-)}{p(x_{n+1}|y^-)}\Big|_{x_{n+1}=1,x^-=0_x^-,y^-=0_y^-}$$
$$=\left(p_x-p_xp_y(l+\sum_t\Delta p_{n+1-t})\right)\log\frac{(1-lp_y)}{1-kp_x-lp_y+p_xp_y(kl+\sum_{s,t}\Delta p_{s-t})} \tag{28}$$
$$=kp_x^2+\frac{k^2p_x^3}{2}-p_x^2p_y(kl+\sum_{s,t}\Delta p_{s-t}+k\sum_t\Delta p_{n+1-t})+O(\Delta t^4),$$

$$p(x_{n+1},x^-,y^-)\log\frac{p(x_{n+1}|x^-,y^-)}{p(x_{n+1}|y^-)}\Big|_{x_{n+1}=0,x^-=1_{x_s},y^-=1_{y_t}}$$
$$=-p_xp_y(1+\Delta p_{s-t})\log\left(1-p_x(1+\Delta p_{n+1-t})\right) \tag{29}$$
$$=p_x^2p_y(1+\Delta p_{s-t}+\Delta p_{n+1-t})\underbrace{+p_x^2p_y\Delta p_{s-t}p_{n+1-t}}_{O(\Delta t^3\Delta p_m^2)}+O(\Delta t^4),$$

$$p(x_{n+1},x^-,y^-)\log\frac{p(x_{n+1}|x^-,y^-)}{p(x_{n+1}|y^-)}\Big|_{x_{n+1}=0,x^-=1_{x_s},y^-=0_y^-}$$
$$=\left(p_x-p_xp_y(l+\sum_t\Delta p_{s-t})\right)\log\frac{1-lp_y}{1-p_x-lp_y+p_xp_y(l+\sum_t\Delta p_{n+1-t})} \tag{30}$$
$$=p_x^2+\frac{p_x^3}{2}-p_x^2p_y(l+\sum_t\Delta p_{n+1-t}+\sum_t\Delta p_{s-t})+O(\Delta t^4),$$

$$p(x_{n+1},x^-,y^-)\log\frac{p(x_{n+1}|x^-,y^-)}{p(x_{n+1}|y^-)}\Big|_{x_{n+1}=0,x^-=0_x^-,y^-=1_{y_t}}$$
$$=\left(p_y-p_xp_y(k+1+\Delta p_{n+1-t}+\sum_s\Delta p_{s-t})\right)\log\frac{1-p_x(k+1+\Delta p_{n+1-t}+\sum_s\Delta p_{s-t})}{(1-p_x(k+\sum_s\Delta p_{s-t}))(1-p_x(1+\Delta p_{n+1-t}))} \tag{31}$$
$$=-p_x^2p_y(k+\sum_s\Delta p_{s-t}+k\Delta p_{n+1-t})\underbrace{-p_x^2p_y\Delta p_{n+1-t}\sum_s\Delta p_{s-t}}_{O(\Delta t^3\Delta p_m^2)}+O(\Delta t^5),$$

$$
\begin{aligned}
&p(x_{n+1},x^-,y^-)\log\frac{p(x_{n+1}|x^-,y^-)}{p(x_{n+1}|y^-)}\bigg|_{x_{n+1}=0,x^-=0_x^-,y^-=0_y^-}\\
&=\left(1-(k+1)p_x-lp_y+p_xp_y(kl+l+\sum_{s,t}\Delta p_{s-t}+\sum_t\Delta p_{n+1-t})\right)\\
&\cdot\log\left[\frac{1-(k+1)p_x-lp_y+p_xp_y(kl+l+\sum_{s,t}\Delta p_{s-t}+\sum_t\Delta p_{n+1-t})}{1-kp_x-lp_y+p_xp_y(kl+\sum_{s,t}\Delta p_{s-t})}\right.\\
&\left.\cdot\frac{1-lp_y}{1-p_x-lp_y+p_xp_y(l+\sum_t\Delta p_{n+1-t})}\right]\\
&=-kp_x^2+p_x^2p_y(kl+k\sum_t\Delta p_{n+1-t}+\sum_{s,t}\Delta p_{s-t})+O(\Delta t^4).
\end{aligned}
\tag{32}
$$

Therefore, combining Eqs. 25 and 27-32, $\mathscr{B}=O(\Delta t^3\Delta p_m^2)$, and thus we can obtain

$$
T_{Y\to X}(k,l;m)=\sum_{i=m}^{m+l-1}I(X,Y;i)+O(\Delta t^3\Delta p_m^2). \tag{33}
$$

Note that we omit higher order terms $O(\Delta t^4)$ in the above derivation.

#### 1.5 Mathematical relation between GC and TE

From Eqs. 8, 14, and 33, we can straightforwardly obtain the following relation between GC and TE

$$
G_{Y\to X}(k,l;m)=2T_{Y\to X}(k,l;m)+O\left(\Delta t^2\Delta p_m^3\right)+O\left(\Delta t^3\Delta p_m^2\right),
$$

where $T_{Y\to X}$ is defined in Eq. 5 with the assumption that $\left\|x_{n+1}^{(k+1)}\right\|_0\le 1$ and $\left\|y_{n+1-m}^{(l)}\right\|_0\le 1$. Next, we will prove that $O\left(\Delta t^3\Delta p_m^2\right)=0$. We collect all the terms with order $O\left(\Delta t^3\Delta p_m^2\right)$ from Eqs. 13, 20, 20, 27, 29, 30, and derive that

$$
\begin{aligned}
O\left(\Delta t^3\Delta p_m^2\right)=&-\frac{1}{\sigma_x^2\sigma_y^2}\left[\mathrm{cov}(x_{n+1},y_{n+1-m}^{(l)})\hat{\mathbf{B}}\,\mathrm{cov}(x_{n+1},y_{n+1-m}^{(l)})^T+\frac{2\sigma_y}{\sigma_x}\mathrm{cov}(x_{n+1},x_n^{(k)})\hat{\mathbf{C}}\,\mathrm{cov}(x_{n+1},y_{n+1-m}^{(l)})^T\right]\\
&-2\left\{p_xp_y^2\sum_t\left(\Delta p_{n+1-t}\sum_{t'\neq t}\Delta p_{n+1-t'}\right)+\frac{1}{2}p_xp_y^2\left[\sum_t\Delta p_{n+1-t}^2-\left(\sum_t\Delta p_{n+1-t}\right)^2\right]\right.\\
&\left.+p_x^2p_y\sum_t\left[\Delta p_{n+1-t}\sum_s\Delta p_{s-t}\right]+p_x^2p_y\left(\sum_{s,t}\Delta p_{s-t}\Delta p_{n+1-t}\right)-p_x^2p_y\sum_t\left[\Delta p_{n+1-t}\sum_s\Delta p_{s-t}\right]\right\}\\
=&-\frac{1}{\sigma_x^2\sigma_y^2}\left[\mathrm{cov}(x_{n+1},y_{n+1-m}^{(l)})\hat{\mathbf{B}}\,\mathrm{cov}(x_{n+1},y_{n+1-m}^{(l)})^T+\frac{2\sigma_y}{\sigma_x}\mathrm{cov}(x_{n+1},x_n^{(k)})\hat{\mathbf{C}}\,\mathrm{cov}(x_{n+1},y_{n+1-m}^{(l)})^T\right]\\
&-p_xp_y^2\left[\left(\sum_{t=n+1-m}^{n+2-m-l}\Delta p_{n+1-t}\right)^2-\sum_{t=n+1-m}^{n+2-m-l}\Delta p_{n+1-t}^2\right]-2p_x^2p_y\left(\sum_{s=n}^{n-k+1}\sum_{t=n+1-m}^{n+2-m-l}\Delta p_{s-t}\Delta p_{n+1-t}\right),
\end{aligned}
\tag{34}
$$

With the assumption that $\left\|x_{n+1}^{(k+1)}\right\|_0\le 1$ and $\left\|y_{n+1-m}^{(l)}\right\|_0\le 1$, components in the first term of Eq. 34 can be rewritten by functions of $p_x$, $p_y$, $\Delta p_{n+1-t}$, and $\Delta p_{s-t}$ as follows

$$\mathrm{cov}(x_{n+1},y^{(l)}_{n+1-m}) = p_x p_y \left[\Delta p_{n+1-(n-m)}, \Delta p_{n+1-(n-m-1)}, \cdots, \Delta p_{n+1-(n-m-l+2)}\right]$$

$$\hat{\mathbf{B}} = \frac{p_y^2}{\sigma_y^2}\left(\mathbf{I}_{l\times l} - 1_{l\times l}\right)$$

$$\mathrm{cov}(x_{n+1},x^{(k)}_{n}) = -p_x^2 1_{1\times k}$$

$$\hat{\mathbf{C}} = \frac{p_x p_y}{\sigma_x\sigma_y}\begin{bmatrix} \Delta p_{(n)-(n-m)} & \Delta p_{(n)-(n-m-1)} & \cdots & \Delta p_{(n)-(n-m-l+2)} \\ \Delta p_{(n-1)-(n-m)} & \Delta p_{(n-1)-(n-m-1)} & \cdots & \Delta p_{(n-1)-(n-m-l+2)} \\ \vdots & \vdots & \ddots & \vdots \\ \Delta p_{(n-k+1)-(n-m)} & \Delta p_{(n-k+1)-(n-m-1)} & \cdots & \Delta p_{(n-k+1)-(n-m-l+2)} \end{bmatrix}.$$

Substituting expressions above into the first term in Eq. 34, we obtain

$$\begin{aligned} O\left(\Delta t^3 \Delta p_m^2\right) =& \frac{p_x^2 p_y^4}{\sigma_x^2\sigma_y^4}\left(\sum_{t=n+1-m}^{n+2-m-l}\sum_{t'=n+1-m}^{n+2-m-l}\Delta p_{n+1-t}\Delta p_{n+1-t'} - \sum_{t=n+1-m}^{n+2-m-l}\Delta p_{n+1-t}^2\right) + 2\frac{p_x^4 p_y^2}{\sigma_x^4\sigma_y^2}\sum_{s=n}^{n-k+1}\sum_{t=n+1-m}^{n+2-m-l}\Delta p_{s-t}\Delta p_{n+1-t} \\ & - p_x p_y^2\left[\left(\sum_{t=n+1-m}^{n+2-m-l}\Delta p_{n+1-t}\right)^2 - \sum_{t=n+1-m}^{n+2-m-l}\Delta p_{n+1-t}^2\right] - 2p_x^2 p_y\left(\sum_{s=n}^{n-k+1}\sum_{t=n+1-m}^{n+2-m-l}\Delta p_{s-t}\Delta p_{n+1-t}\right) \\ =& 0. \end{aligned}$$

Note that we omit higher order terms $O(\Delta t^4)$ in the above derivation. Therefore, we prove the Theorem 4 that

$$G_{Y\to X}(k,l;m) = 2T_{Y\to X}(k,l;m) + O\left(\Delta t^2 \Delta p_m^3\right). \quad (35)$$

### 2 Mechanism underlying successful network reconstruction using pairwise causal inference

Here we demonstrate the validity of pairwise inference on pulse-output signals in the reconstruction of network structural connectivity. It has been noticed that pairwise causal inference may potentially fail to distinguish the direct interactions from the indirect ones in a network. For example, in the case that $Y \to W \to X$ where "$\to$" denotes a direct connection, the indirect interaction from Y to X may possibly be mis-inferred as a direct interaction via pairwise causality measures especially when the activity signals are continuous-valued as shown in Fig. 11B. However, this type of mistake does not happen in our case of pulse-output signals as explained below. Here we take TDCC as an example to explain the underlying reason of successful reconstruction.

Denote

$$\delta p_{Y\to X} = p(x_n = 1|y_{n-m} = 1) - p(x_n = 1|y_{n-m} = 0)$$

as the increment of probability of generating a pulse output by node $X$ at time step $n$ induced by a pulse-output signal of node $Y$ at an earlier time step $n-m$. From Eq. 6, we have

$$C(X,Y;m) = \delta p_{Y\to X}\sqrt{\frac{p_y - p_y^2}{p_x - p_x^2}}. \tag{36}$$

Denote $S_1$ and $S_2$ as the coupling strength from node $Y$ to node $W$ and from node $W$ to node $X$, respectively. Then the increment $\delta p_{Y\to X}$ is a function of $S_1$ and $S_2$ and the Taylor expansion of $\delta p_{Y\to X}$ with respect to $S_1$ and $S_2$ has the following form

$$\delta p_{Y\to X} = \alpha_0 + \alpha_1 S_1 + \alpha_2 S_2 + \alpha_3 S_1^2 + \alpha_4 S_1 S_2 + \alpha_5 S_2^2 + o(S_1 S_2), \tag{37}$$

where the symbol "o" stands for higher order terms. If $S_1 = 0$ or $S_2 = 0$, then the nodes $X$ and $Y$ are independent from the connection structure, *i.e.*,

$$\delta p_{Y\to X}\Big|_{S_1=0} = 0 \quad \text{and} \quad \delta p_{Y\to X}\Big|_{S_2=0} = 0.$$

Therefore, we have $\alpha_0 = \alpha_1 = \alpha_2 = \alpha_3 = \alpha_5 = 0$ in Eq. 37 and $\delta p_{Y\to X} = \alpha_4 S_1 S_2 + o(S_1 S_2)$. Similarly, the Taylor expansion of $\delta p_{Y\to W}$ and $\delta p_{W\to X}$ with respect to $S_1$ and $S_2$ have the form

$$\delta p_{Y\to W} = \beta_1 S_1 + O(S_1^2) \quad \text{and} \quad \delta p_{W\to X} = \beta_2 S_2 + O(S_2^2),$$

which are numerically verified in Fig. 8C. Thus, we have

$$\delta p_{Y\to X} = O(\delta p_{Y\to W} \cdot \delta p_{W\to X}), \tag{38}$$

as shown in Fig. 8A-B (bottom right inset) for examples of 3-neuron HH networks. From Eqs. 36 and 38, we have

$$C(X,Y;m) = O\left(C(W,Y;m)\cdot C(X,W;m)\right)$$

as shown in Fig. 8A-B. Because the influence of a single input pulse signal is often small (*e.g.*, in the HH neural network with physiologically realistic coupling strengths corresponding to excitatory postsynaptic potential less than 1 mV, the absolute value of the increment $|\delta p|$ is less than 0.01 measured from simulation, as shown in Fig. 8C), the causal value $C(X,Y;m)$ from indirect interaction will be significantly smaller than $C(W,Y;m)$ or $C(X,W;m)$ from the direct interaction. Therefore, direct connections and indirect connections are distinguishable when performing pairwise inference on pulse-output signals.

Furthermore, we also shows the relation between $\delta p_{Y\to X}$ and $\Delta p_m$, which is introduced in derivations of our theorems.

$$\begin{aligned}
\delta p_{Y\to X} &= \frac{p\left(x_n=1, y_{n-m}=1\right)}{p\left(y_{n-m}=1\right)} - \frac{p\left(x_n=1, y_{n-m}=0\right)}{p\left(y_{n-m}=0\right)} \\
&= \frac{p\left(x_n=1, y_{n-m}=1\right)}{p\left(y_{n-m}=1\right)} + \frac{p\left(x_n=1, y_{n-m}=1\right)}{p\left(y_{n-m}=0\right)} - \frac{p\left(x_n=1, y_{n-m}=1\right)}{p\left(y_{n-m}=0\right)} - \frac{p\left(x_n=1, y_{n-m}=0\right)}{p\left(y_{n-m}=0\right)} \\
&= \left[\frac{p\left(x_n=1, y_{n-m}=1\right)}{p\left(x_n=1\right)p\left(y_{n-m}=1\right)} - 1\right]\cdot\frac{p\left(x_n=1\right)}{p\left(y_{n-m}=0\right)} \\
&= \Delta p_m \cdot \frac{p_x \Delta t}{1 - p_y \Delta t} \\
&\approx \Delta p_m \cdot p_x \Delta t
\end{aligned} \tag{39}$$

We've shown that $\delta p_{Y\to X}$ is proportional to $S$, shown in Fig. 8C, and $\Delta p_m$ is insensitive to $\Delta t$, as shown in Fig. 2. Therefore, $\Delta p_m$ is asymptotically proportional to $O(S)$, and $\delta p_{Y\to X}$ is asymptotically proportional to $O(S\cdot\Delta t)$,

$$\Delta p_m \propto O(S), \qquad \delta p_{Y\to X} \propto O(S\cdot\Delta t). \tag{40}$$

### 3 Detailed HH model

#### 3.1 Hodgkin-Huxley (HH) neural network model of only excitatory population

The dynamics of the $i$th neuron of an HH network is governed by

$$C\frac{dV_i}{dt} = -G_{\mathrm{Na}}m_i^3h_i(V_i - V_{\mathrm{Na}}) - G_{\mathrm{K}}n_i^4(V_i - V_{\mathrm{K}}) - G_L(V_i - V_L) + I_i^{\mathrm{input}}, \tag{41}$$

$$\frac{dz_i}{dt} = (1-z_i)\alpha_z(V_i) - z_i\beta_z(V_i), \quad \text{for } z = m,h,n, \tag{42}$$

where $C$ is the cell membrane capacitance; $V_i$ is the membrane potential (voltage); $m_i$, $h_i$, and $n_i$ are gating variables; $V_{\mathrm{Na}}, V_{\mathrm{K}}$, and $V_L$ are the reversal potentials for the sodium, potassium, and leak currents, respectively; and $G_{\mathrm{Na}}, G_{\mathrm{K}}$, and $G_L$ are the corresponding maximum conductances. The rate variables $\alpha_z$ and $\beta_z$ are defined as Dayan and Abbott (2001)

$$\begin{aligned}
\alpha_m(V) &= \frac{0.1V+4}{1-\exp(-0.1V-4)}, & \beta_m(V) &= 4\exp\left(\frac{-(V+65)}{18}\right), \\
\alpha_h(V) &= 0.07\exp\left(\frac{-(V+65)}{20}\right), & \beta_h(V) &= \frac{1}{1+\exp(-3.5-0.1V)}, \\
\alpha_n(V) &= \frac{0.01V+0.55}{1-\exp(-0.1V-5.5)}, & \beta_n(V) &= 0.125\exp\left(\frac{-(V+65)}{80}\right).
\end{aligned}$$

The input current $I_i^{\mathrm{input}}$ has the form $I_i^{\mathrm{input}} = -G_i(t)(V_i - V_E)$, with $V_E$ being the excitatory reversal potential. The conductance $G_i(t)$ is defined as

$$G_i(t) = f\sum_l H(t - s_{il}) + \sum_j A_{ij}S\sum_l H(t-\tau_{jl}),$$

with $s_{il}$ being the $l$th spike time of the external Poisson input with strength $f$ and rate $\nu$. The spike-induced conductance change $H(t)$ is defined by Dayan and Abbott (2001)

$$H(t) = \frac{\sigma_d\sigma_r}{\sigma_d - \sigma_r}\left[\exp\left(-\frac{t}{\sigma_d}\right) - \exp\left(-\frac{t}{\sigma_r}\right)\right]\Theta(t), \tag{43}$$

where $\sigma_d$ and $\sigma_r$ are the decay and rise time scale, respectively, and $\Theta(\cdot)$ is the Heaviside function. $\mathbf{A} = (A_{ij})$ is the adjacency matrix with $A_{ij} = 1$ indicating a direct connection from neuron $j$ to neuron $i$ and $A_{ij} = 0$ indicating no connection from neuron $j$ to neuron $i$, $S$ is the coupling strength, and $\tau_{jl}$ is the $l$th spike time of the $j$th neuron.

We take the parameters as in Ref. Dayan and Abbott (2001) that $C = 1\,\mu\mathrm{F}\cdot\mathrm{cm}^{-2}$, $V_{\mathrm{Na}} = 50$ mV, $V_{\mathrm{K}} = -77$ mV, $V_L = -54.387$ mV, $G_{\mathrm{Na}} = 120$ mS·cm$^{-2}$, $G_{\mathrm{K}} = 36$ mS·cm$^{-2}$, $G_L = 0.3$ mS·cm$^{-2}$, and $V_E = 0$ mV. We set synaptic time constants as $\sigma_r = 0.5$ ms and $\sigma_d = 3.0$ ms. For simplicity, we set the Poisson input parameters as $f = 0.1$ mS·cm$^{-2}$ and $\nu = 100$ Hz, unless indicated otherwise. However, the conclusions shown in this work hold for a wide range of parameters corresponding to different dynamical regimes.

When the voltage $V_i$ reaches the firing threshold, $V_{\mathrm{th}} = -50$ mV, we say the $i$th neuron generates a spike at this time. Instantaneously, all of its postsynaptic neurons receive this spike and the affected change of conductance follows Eq. 43.

#### 3.2 HH neural network model of both excitatory and inhibitory populations

For the HH network consisting of both excitatory and inhibitory neurons, the dynamics of the $i$th HH neuron is also governed by Eqs. 41 and 42. But the input current $I_i^{\mathrm{input}}$ is given by

$$I_i^{\mathrm{input}} = -G_i^E(t)(V_i - V_E) - G_i^I(t)(V_i - V_I),$$

where $G_i^E(t)$ and $G_i^I(t)$ are excitatory and inhibitory conductances, respectively, $V_E$ and $V_I$ are the corresponding reversal potentials. The conductances are defined as

$$G_i^E(t) = f\sum_l H(t - s_{il}; \sigma_d^E, \sigma_r^E) + \sum_j A_{ij} S^E \sum_l H(t - \tau_{jl}; \sigma_d^E, \sigma_r^E),$$
$$G_i^I(t) = \sum_j A_{ij} S^I \sum_l H(t - \tau_{jl}; \sigma_d^I, \sigma_r^I),$$

where $H(\cdot)$ is given in Eq. 43 with parameters $\sigma_d^E$ ($\sigma_d^I$) and $\sigma_r^E$ ($\sigma_r^I$) being the decay and rise time scale of excitation (inhibition); $S^E$ and $S^I$ are the excitatory and inhibitory coupling strengths, respectively. The parameters are set as $V_E = 0$ mV, $V_I = -80$ mV, $\sigma_r^E = 0.5$ ms, $\sigma_d^E = 3.0$ ms, $\sigma_r^I = 0.5$ ms, $\sigma_d^I = 7.0$ ms. The HH neural network here and the previous one with only excitatory population are efficiently simulated by an adaptive exponential time differencing algorithm introduced in Ref. Tian and Zhou (2020).

### Properties of pulse-output signals

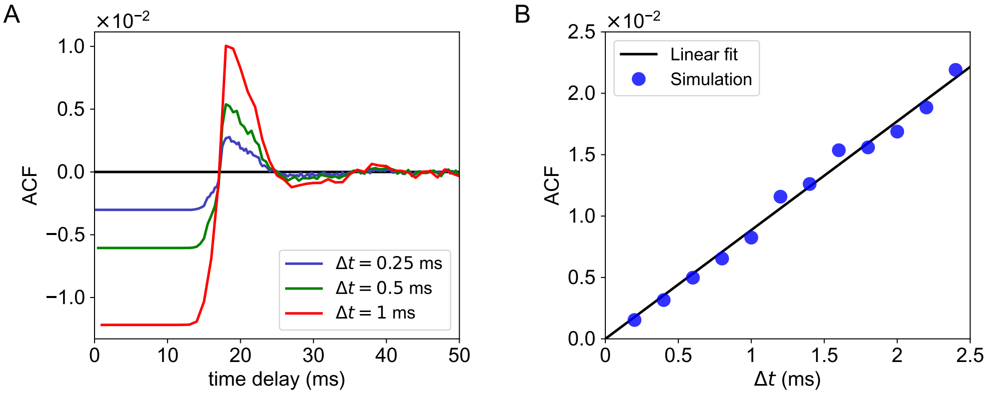


Figure 1: Relation between ACF and sampling time bin $\Delta t$ of pulse-output signals. ($A$) ACF curves as a function of time delay with $\Delta t = 0.25$ (blue), 0.5 (green), and 1 ms (red), respectively. Note that ACF with 0 time delay is not plotted. ($B$), ACF values at a fixed time delay 20 ms plotted as a function of $\Delta t$. The black line is a linear fit with $R^2 = 0.985$ which is consistent with the derivation in Eq. 10. When $\Delta t$ is sufficiently small, the magnitude of auto-correlation of binary time series is also small, indicating that the binary time series become almost whitened.

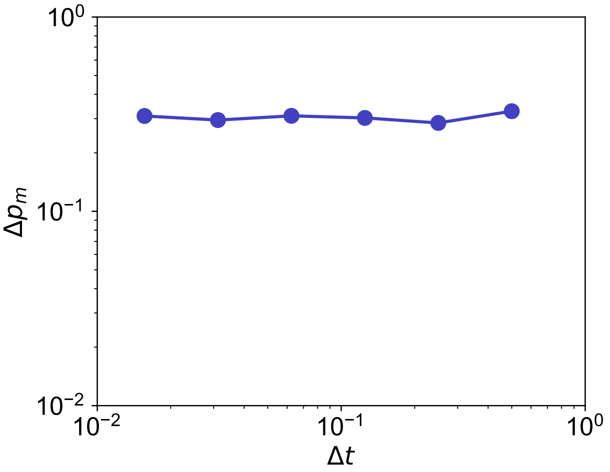


Figure 2: $\Delta p_m$ is insensitive to sampling resolution $\Delta t$. The result is obtained from neuron $Y$ to neuron $X$ in an HH network of 10 excitatory neurons. The HH network is randomly connected with connection probability 0.25, and there is a unidirectional connection from $Y$ to $X$ with coupling strength $S$. The parameters are set as a fixed time delay 3 ms and $S = 0.02$ mS·cm$^{-2}$.

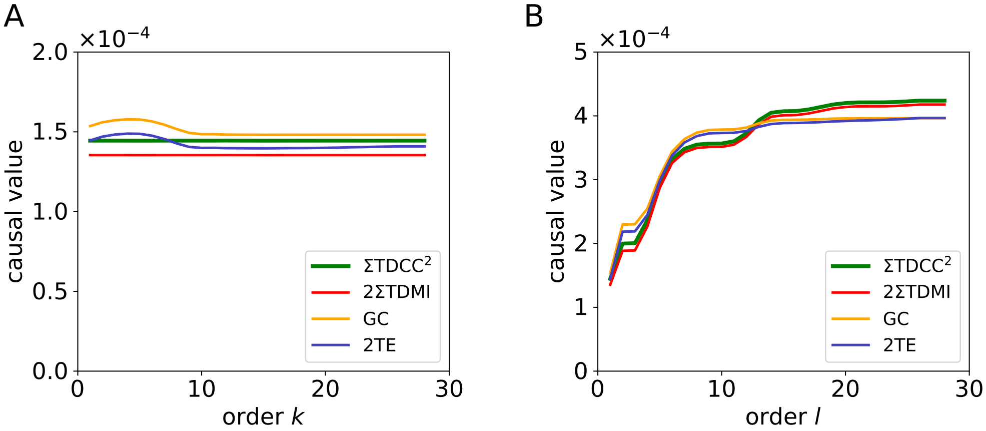


Figure 3: Causal values as a function of ($A$) order $k$ and ($B$) order $l$ which are computed from neuron $Y$ and neuron $X$ in the same HH network as in Fig. 2. A large $\Delta t = 1.5$ms is applied in the computation of causal values. Other parameters are set as $m = 2$ (time delay is 3 ms), $f = 0.1$ mS$\cdot$cm$^{-2}$, $\nu = 300$ Hz, $S = 0.02$ mS$\cdot$cm$^{-2}$, and $l = 1$ in ($A$) and $k = 1$ in ($B$). The causal values in both ($A$) and ($B$) are all significantly greater than those of randomly surrogate time series with the p-value $p < 0.05$. The relations among the four causality measures revealed by Theorems 1-4 in the main text still holds when choosing the orders of $k = 28$ in ($A$) or $l = 28$ in ($B$), in both cases the event $\left\|x_{n+1}^{(k+1)}\right\|_0 \geq 2$ or $\left\|y_{n+1-m}^{(l)}\right\|_0 \geq 2$ occurs with a frequency more than 44%. This result indicates that the assumption of $\left\|x_{n+1}^{(k+1)}\right\|_0 \leq 1$ and $\left\|y_{n+1-m}^{(l)}\right\|_0 \leq 1$ is a sufficient but not necessary condition in the derivation of the quantitative relation between TE and TDMI.

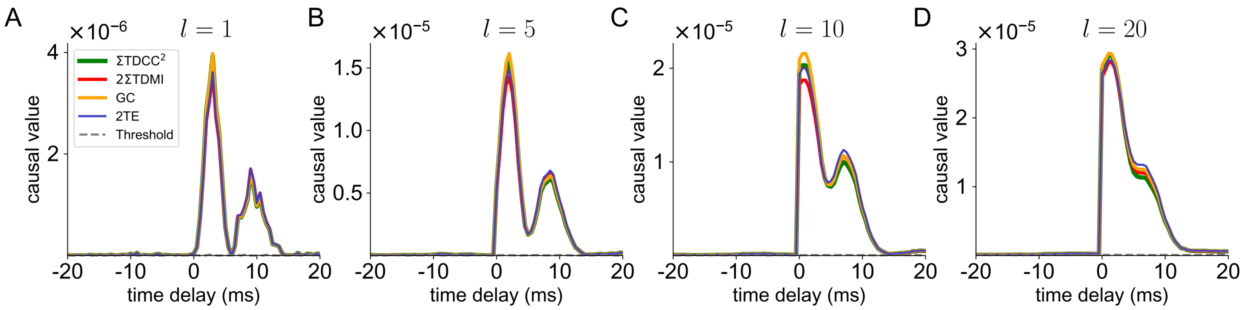


Figure 4: Dependence of causal values on the parameter of time delay, with different choice of order $l$. In (A)-(D), order $l$ is equal to 1, 5, 10 and 20, respectively. The gray dashed curve is the significance level of causality for unconnected pairs. Data in (A) are the same as in Fig. 3C in the main text and are reproduced here for comparison with other cases. Note that there is a well-separated second peak in (A) around 9.5 ms, which results from the incomplete estimation of causal information due to the choice of small value of $l$ (*e.g.*, $l = 1$). The second peak gradually disappears as $l$ increases. On the one hand, for different choice of $l$, the mathematical relations in Theorem 1-4 in the main text always hold. On the other hand, order $l = 1$ is sufficient for the inference of a correct direction of causal connection, since the causal values of direct connections is significantly distinguishable from pairs without direct connections. The colors and other parameters are set the same as those in Fig. 3C in the main text.

### Consistency among causality measures across different dynamical regimes

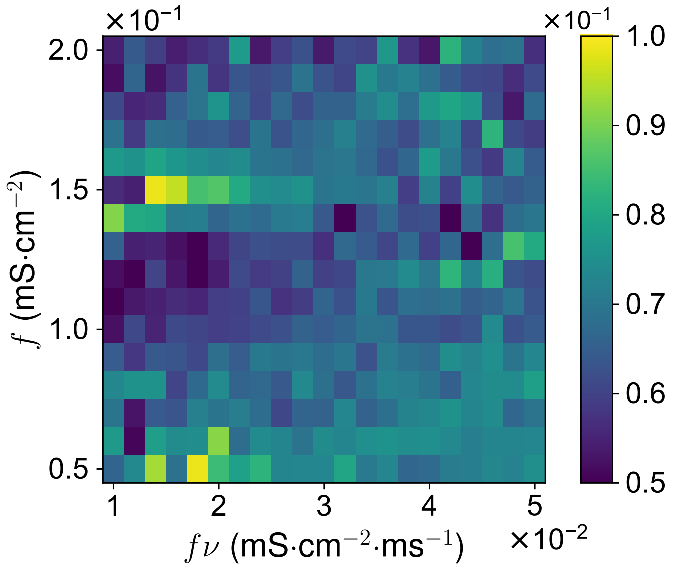


Figure 5: Relative error of causal values with different external Poisson input parameters $f$ and $\nu$. The result is obtained from neuron $Y$ to neuron $X$ in the same HH network in Fig. 2. Here, the relative error is computed by $\frac{\max\{\Sigma\mathrm{TDCC}^2, 2\Sigma\mathrm{TDMI}, \mathrm{GC}, \mathrm{TE}\} - \min\{\Sigma\mathrm{TDCC}^2, 2\Sigma\mathrm{TDMI}, \mathrm{GC}, \mathrm{TE}\}}{\max\{\Sigma\mathrm{TDCC}^2, 2\Sigma\mathrm{TDMI}, \mathrm{GC}, \mathrm{TE}\}}$ and small relative error indicates that mathematical relations revealed by Theorems 1-4 in the main text hold for a wide range of Poisson input parameters. Other parameters are set as $\Delta t = 0.5\,\mathrm{ms}$, $k = l = 1$, $S = 0.01$ mS·cm$^{-2}$, and $m = 6$ (time delay is 3 ms).

### Reconstruction of structure connectivity for asynchronous state

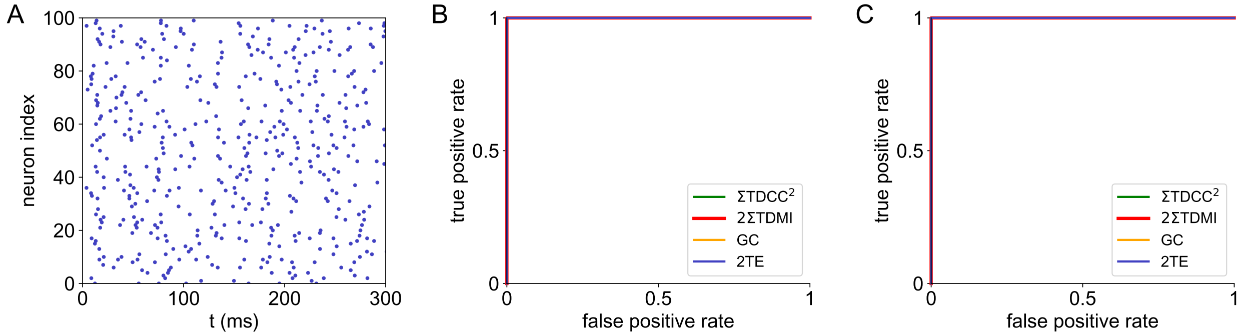


Figure 6: Performance of the causality measures in an HH network in the asynchronous state. The network is composed of 100 excitatory neurons randomly connected with probability 0.25. (*A*) Raster plot of neuronal firing indicating that the network is in an asynchronous state. (*B*) ROC curves of the full HH network with AUC $= 1$. (*C*) ROC curves of an HH subnetwork of 20 neurons with AUC $= 1$. The green curve represents the summation of squared TDCC $C(X,Y;m)$, the red curve represents twice of the summation of TDMI $I(X,Y;m)$, the orange curve stands for GC $G_{Y\to X}(k,l;m)$, and the blue curve stands for twice of TE $T_{Y\to X}(k,l;m)$. The ROC curves for TDCC, TDMI, GC, and TE overlap with each other. The parameters are set as $\Delta t = 0.5\,\mathrm{ms}$, $k = l = 1$, $S = 0.02$ mS$\cdot$cm$^{-2}$, and $m = 6$ (time delay is 3 ms).

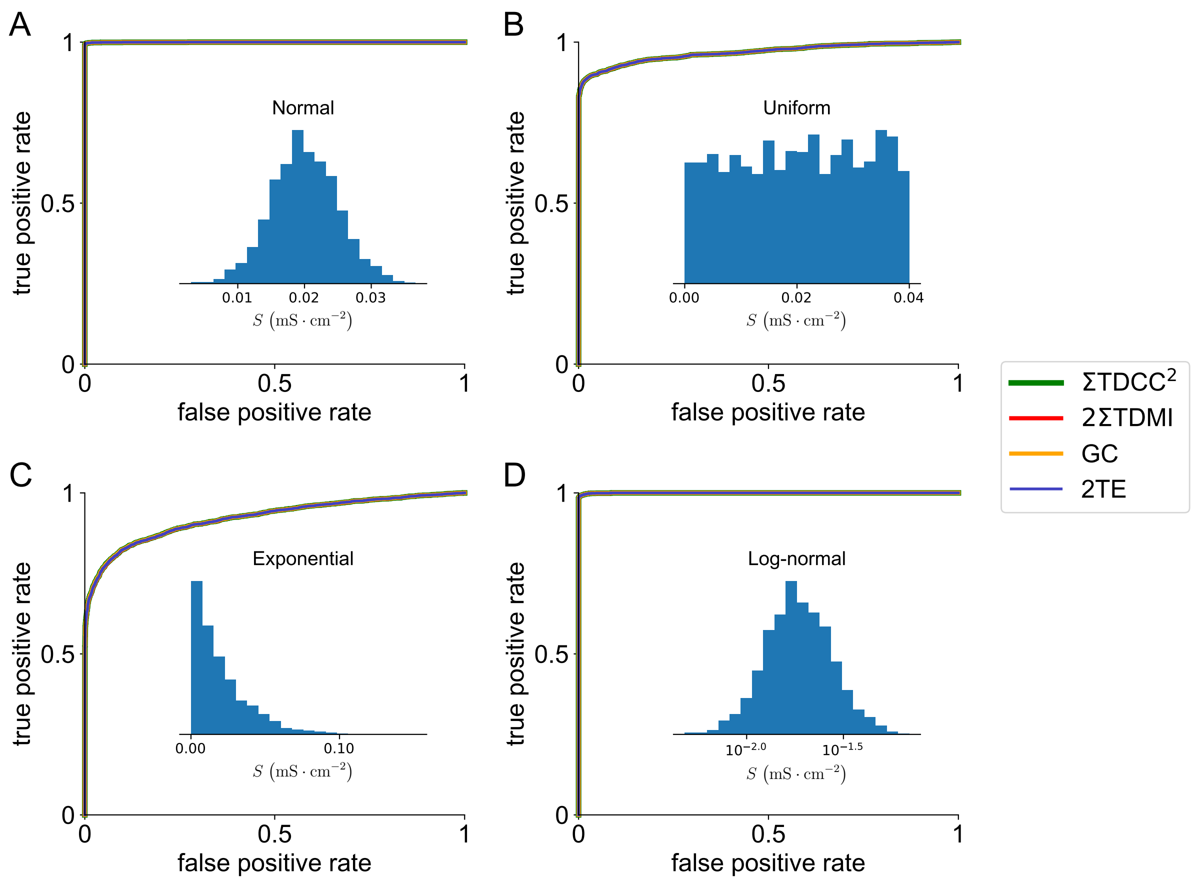


Figure 7: Performance of causality measures in HH networks with heterogeneous structural connectivity. The networks are composed of 100 excitatory neurons with the entry $A_{ij}$ in the adjacency matrix following the Bernoulli distribution (probability of 0.25 being 1). For those connected pairs, *e.g.*, $A_{ij} = 1$, the corresponding coupling strength from neuron $j$ to neuron $i$ is sampled from various distributions: (*A*) normal, (*B*) uniform, (*C*) exponential, and (*D*) log-normal distributions. (*A*-*D*) The AUC values of HH networks are 1.0, 0.97, 0.92, and 1.0, respectively. Note that all ROC curves are virtually overlap with each other, which again is consistent with Theorems 1-4 in the main text. The colors are the same as those in Fig. 6. Inset: The corresponding histograms of the coupling strength, $S$, among all connected pairs in networks. The parameters are set as $\Delta t = 0.5\,\text{ms}$, $k = l = 1$.

### Dependence of $\delta p$ on $S$

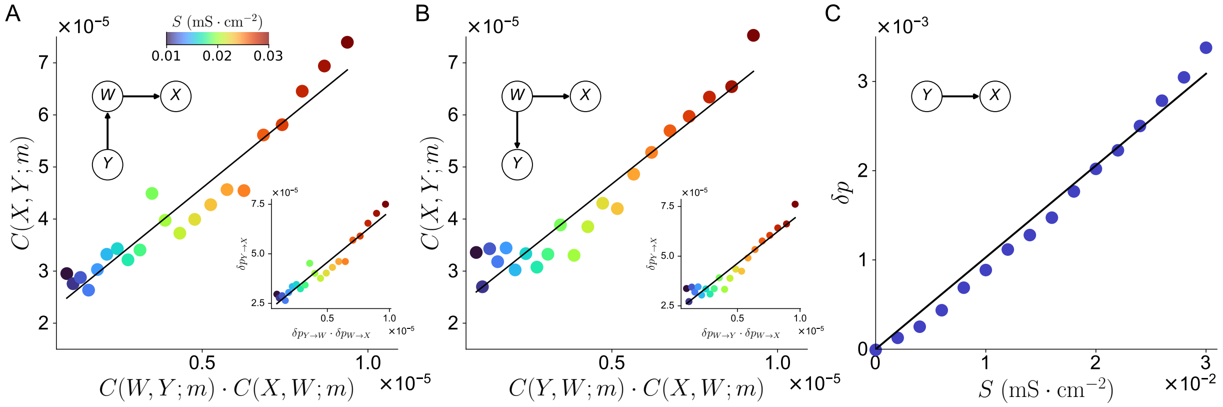


Figure 8: The relations of TDCC between indirectly coupled pair and directly connected pairs, and the dependence of the increment $\delta p$ of directly connected pair on the coupling strength $S$. *(A-B)*: $C(X,Y,m)$ (of indirectly coupled pair) is linearly correlated with the product of directly connected pairs $C(W,Y;m)$ and $C(X,W;m)$ for a 3-neuron HH network with structural connectivity given in the inset (top left) in *(A)*. $C(X,Y,m)$ (of indirectly coupled pair) is linearly correlated with the product of directly connected pairs $C(Y,W;m)$ and $C(X,W;m)$ for a 3-neuron network with structural connectivity given in the inset (top left) in *(B)*. The black line is a linear fit with $R^2 = 0.928$ in *(A)* and $R^2 = 0.913$ in *(B)*. The Inset (bottom right) in *(A)*: $\delta p_{Y \to X}$ (of indirectly coupled pair) is linearly correlated with the product of directly connected pairs $\delta p_{Y \to W}$ and $\delta p_{W \to X}$. The black line is a linear fit with $R^2 = 0.930$. The Inset (bottom right) in *(B)*: $\delta p_{Y \to X}$ (of indirectly coupled pair) is linearly correlated with the product of directly connected pairs $\delta p_{W \to Y}$ and $\delta p_{W \to X}$. The black line is a linear fit with $R^2 = 0.917$. *(C)* $\delta p_{Y \to X}$ is proportional to the coupling strength $S$ in a 2-neuron HH network with structural connectivity given in the inset (top left). The black line is a linear fit with $R^2 = 0.992$. The colormap in *(A-B)* (including insets) indicates the magnitude of coupling strength $S$ defined by the colorbar in *(A)*. The parameters are set as $\Delta t = 0.5$ ms, and $m = 6$ (time delay is 3 ms).

### Reconstruction of structure connectivity with experimental data

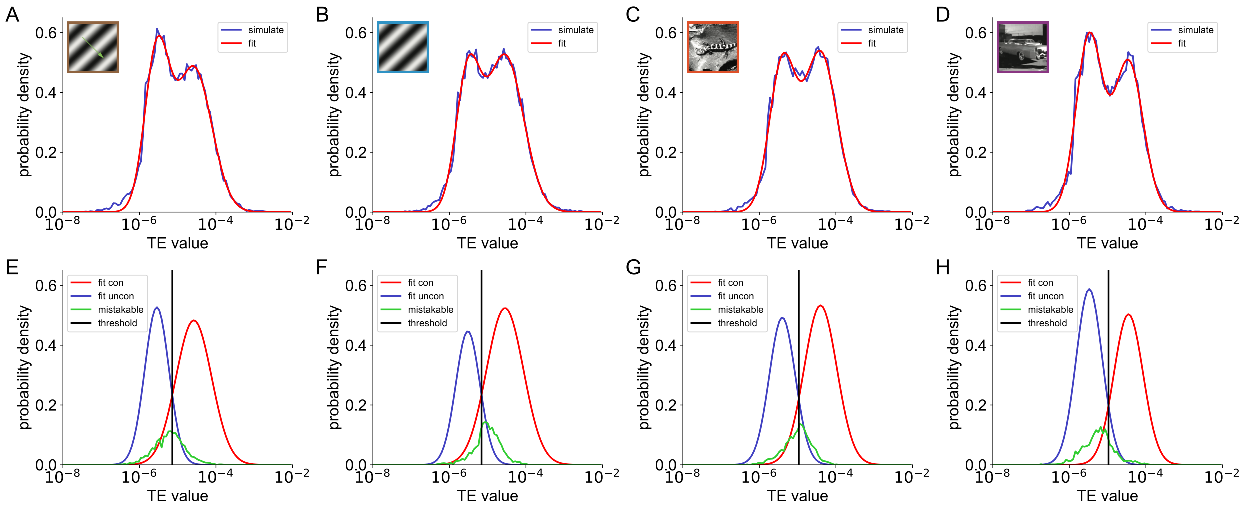


Figure 9: Reconstruction of structural connectivity by the assumption of log-normal distribution of causal values for experimental spike data. (Top panel): Distribution of TE values in the network composed by the observed neurons in experiment under visual stimuli of (*A*) drifting gratings, (*B*) static gratings, (*C*) natural scenes, and (*D*) natural movie. The blue and red curves are the computed and fitted distributions, respectively. (Bottom panel): Distributions of fitted TE values from connected (red) and unconnected (blue) pairs which are obtained from the fitting in top panel. The black vertical line is the optimal inference threshold and the green curve is the mistakable causal values. We use the experimental spike data (sections id 715093703 at `https://allensdk.readthedocs.io/`) with signal-to-noise ratio greater than 4 and firing rate greater than 0.08 Hz. The parameters are set as $k = 1, l = 5, \Delta t = 1$ ms, and $m = 1$ (time delay is 1 ms).

### Verify the log-normal distributed assumption for causality measures

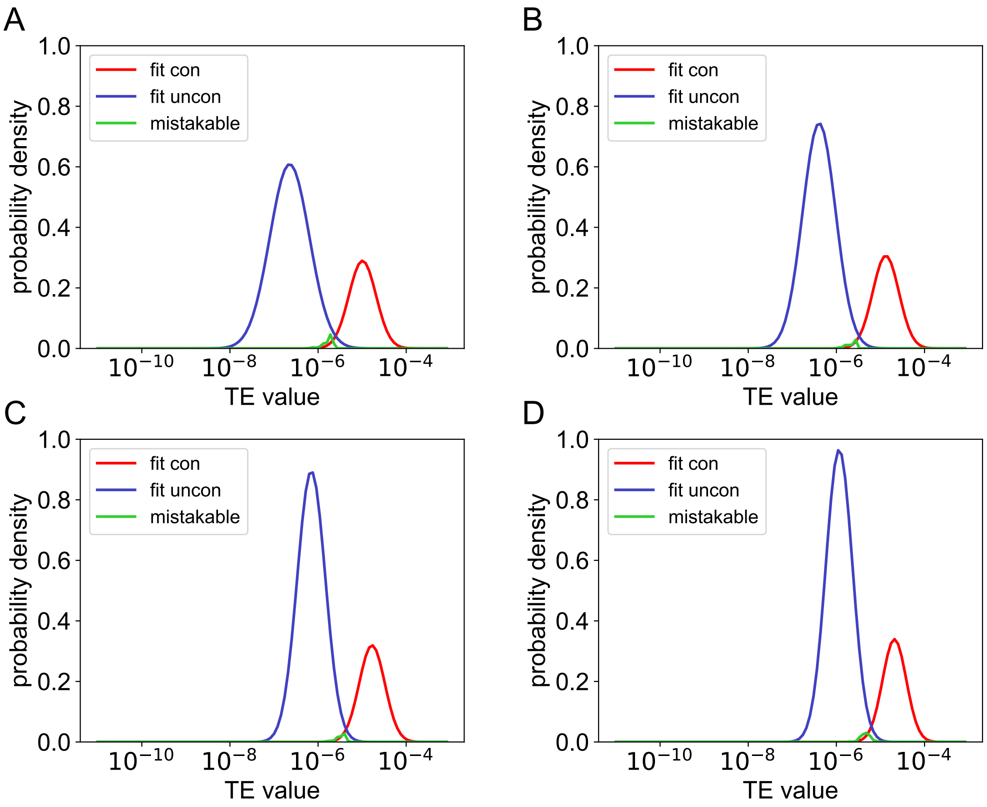


Figure 10: Reconstruction of structural connectivity by the assumption of log-normal distribution of causal values for an HH network of 100 excitatory neurons. The entry $A_{ij}$ in the adjacency matrix follows a Bernoulli distribution with probability of 0.25 being 1. For the connected pairs, *e.g.*, $A_{ij} = 1$, the corresponding coupling strength from neuron $j$ to neuron $i$ is sampled from a log-normal distribution. The parameters are the same as those in Fig. 6 except that the Poisson input rate is $\nu = 90$ Hz in (*A*), $\nu = 100$ Hz in (*B*), $\nu = 110$ Hz in (*C*), and $\nu = 120$ Hz in (*D*). The colors are the same as those in Fig. 9.

### Continuous-valued signals breaks the mathematical relations among four causality measures

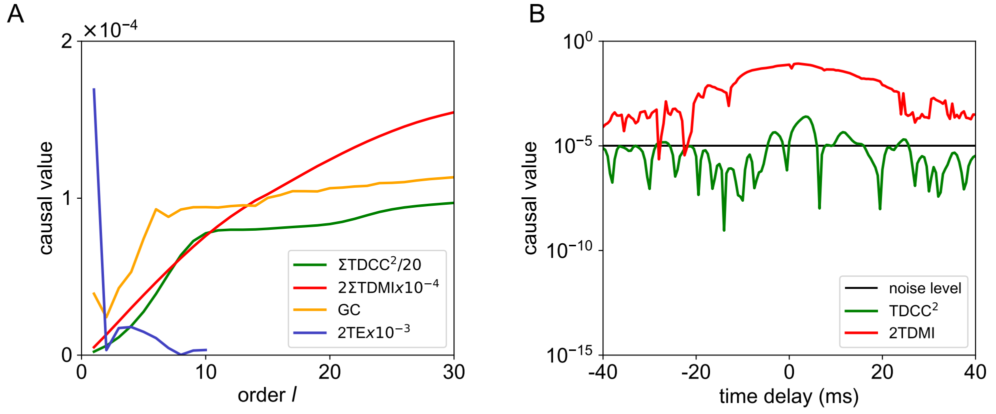


Figure 11: The mathematical relations among the causality measures in Theorems 1-4 in the main text do not hold for continuous-valued voltage time series in the same HH network in Fig. 3. (*A*) TDCC, TDMI, GC, and TE as a function of order $l$ computed from continuous-valued voltage time series. The order $l$ for TE is cut off at $l = 10$ due to the exponential increase of data requirement. (*B*) TDCC and TDMI as a function of time delay with positive (negative) delay corresponding to the calculation of causal values from $Y$ to $X$ (from $X$ to $Y$). The black line represents the noise level, which is obtained as the largest value of TDCC (TDMI) after shuffling the time series and computing TDCC (TDMI) between the shuffled signals for 100 times. A bidirectional connection between $X$ and $Y$ will be incorrectly inferred by TDMI due to the strong self-correlation of the continuous-valued voltage time series. The parameters are set as order $k = l$ and $m = 1$ (time delay is 0.5 ms) in (*A*), and $S = 0.02$ mS$\cdot$cm$^{-2}$, $\Delta t = 0.5$ ms in (*A*) and (*B*).

### Reconstruction of structure connectivity in more general situations

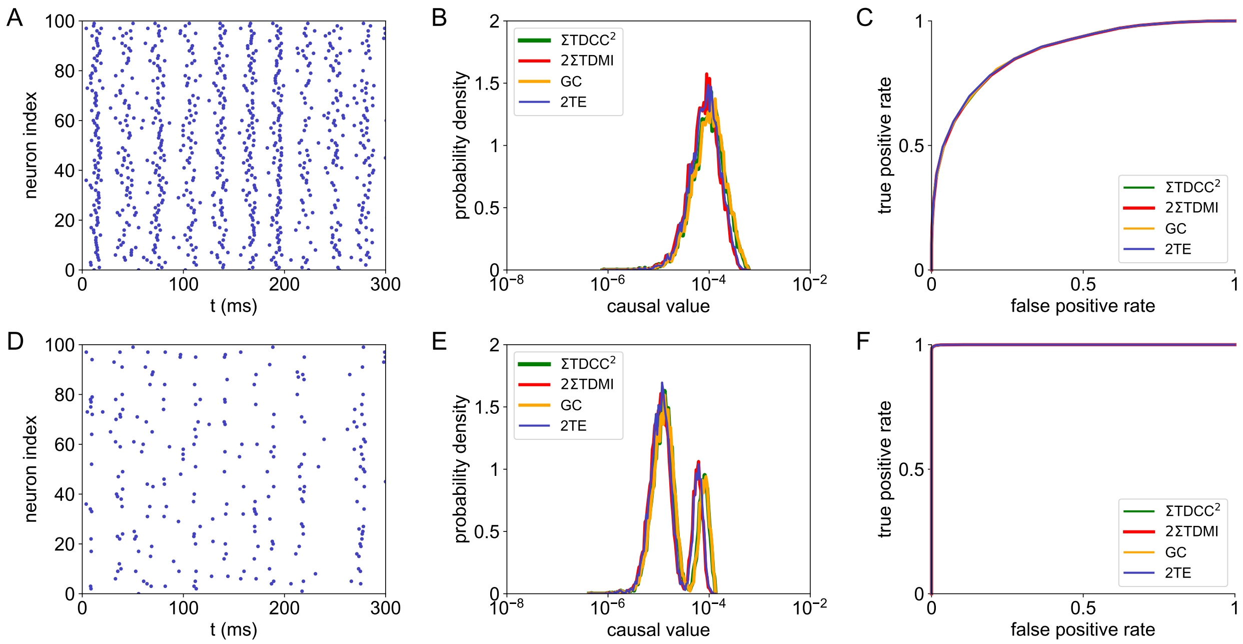


Figure 12: Performance of the causality measures in an HH network of 100 excitatory neurons in the nearly synchronous state. (Top panel): Results using the original spike train. (Bottom panel): Results using the spike train from desynchronized sampling that only samples the pulse-output signals in asynchronous time intervals. (*A*,*D*): Raster plot of the neuronal firing. (*B*, *E*): The distribution of causal values of each pair of neurons in the whole network. (*C*, *F*): ROC curves of the HH network with AUC = 0.88 (upper) and AUC = 0.99 (lower). The ROC curves for TDCC, TDMI, GC, and TE nearly overlap. The colors and parameters are the same as those in Fig. 6, except that the coupling strength $S = 0.028$ mS·cm$^{-2}$.

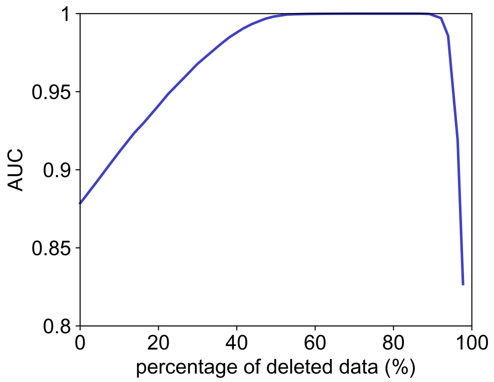


Figure 13: AUC as a function of percentage of deleted data in the spike train of the HH network in Fig. 12A. 78 % of the spike data are deleted by performing desynchronized sampling (*i.e.*, only spike data in asynchronous time intervals are kept) in Fig. 12D.

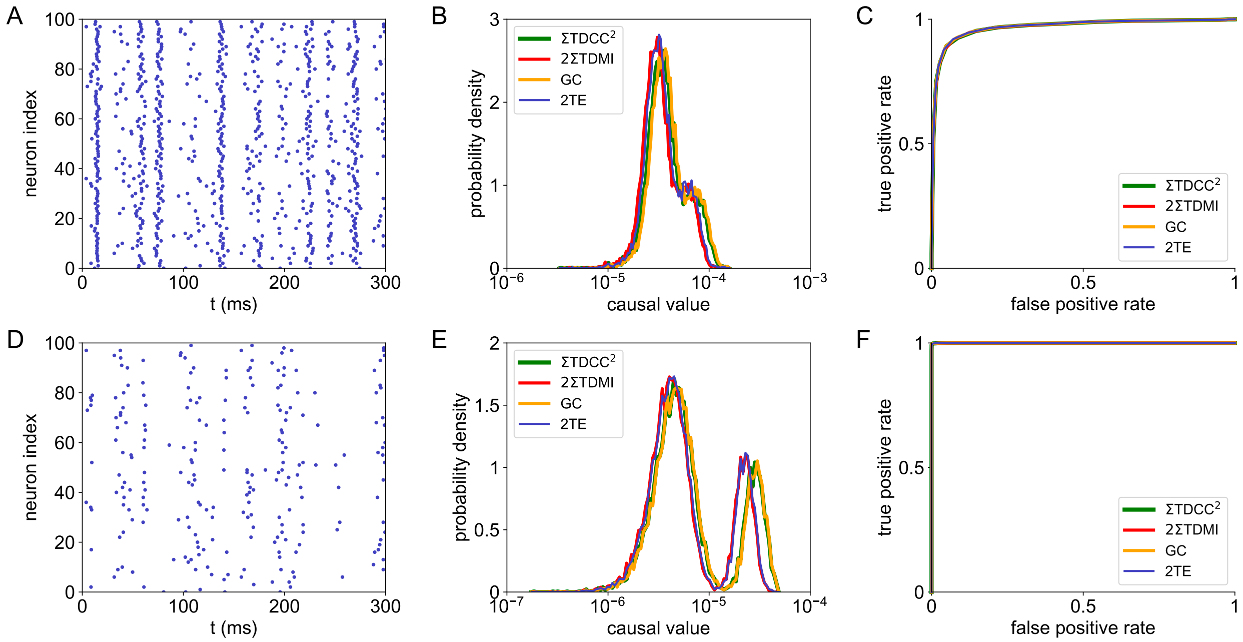


Figure 14: Performance of the causality measures in an HH network of 100 excitatory neurons receiving correlated external Poisson inputs. The correlation coefficient of the Poisson input to each neuron is 0.35. (Top panel): Results using the original spike train. (Bottom panel): Results using the spike train from desynchronized sampling. (*A*, *D*): Raster plot of the neuronal firing. (*B*, *E*): The distribution of causal values of each pair of neurons in the whole network. (*C*, *F*): ROC curves of the HH network with AUC $= 0.96$ (upper) and AUC $= 0.99$ (lower). The ROC curves for TDCC, TDMI, GC, and TE nearly overlap. The colors and parameters are the same as those in Fig. 6.

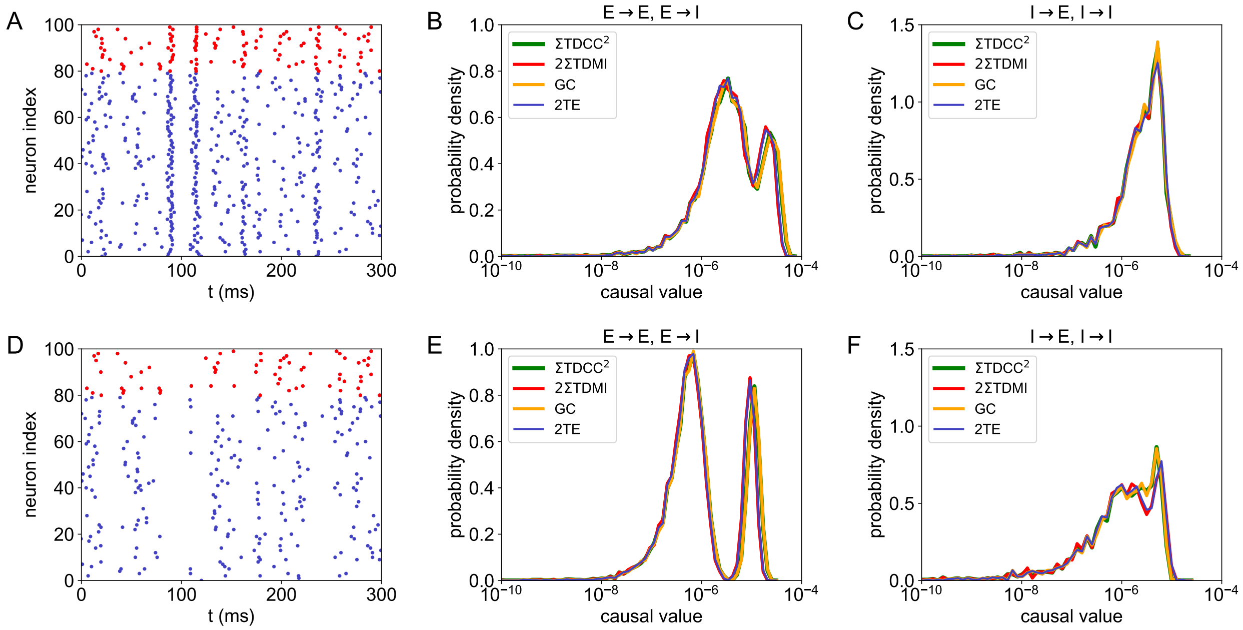


Figure 15: Performance of the causality measures in an HH network of 80 excitatory and 20 inhibitory neurons. The neurons are randomly connected with probability 0.25. (Top panel): Results using the original spike train. (Bottom panel): Results using the spike train from desynchronized sampling. (*A, D*) Raster plot of the neuronal firing. The blue and red dots indicate the excitatory and inhibitory neurons, respectively. (*B, E*) The distribution of causal values of each pair of neurons with the presynaptic neuron being excitatory. (*C, F*) The distribution of causal values of each pair of neurons with the presynaptic neuron being inhibitory. The colors and parameters are the same as those in Fig. 6. The AUC values for (B, C, E, F) are 0.96, 0.71, 1, and 0.99, respectively. The coupling strength is $S^E = 0.02$ mS·cm$^{-2}$ and $S^I = 0.08$ mS·cm$^{-2}$. The correlation coefficient of the Poisson input to each neuron is 0.15.

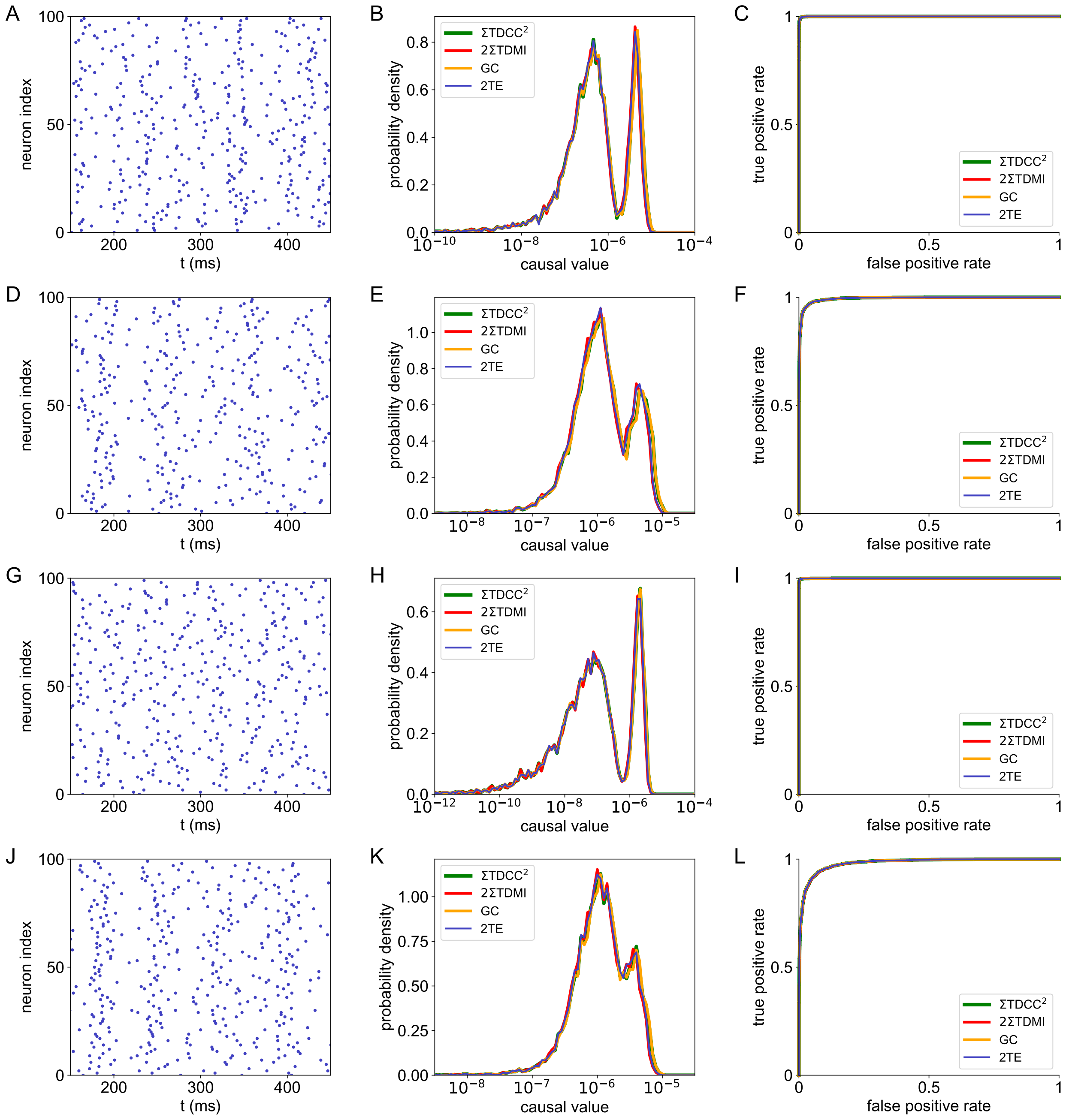

Figure 16: Performance of the causality measures in (*A-C*) I&F, (*D-F*) Izhikevich, (*G-I*) FitzHugh-Nagumo, (*J-L*) Morris-Lecar network of 100 excitotary neurons randomly connected with probability 0.25. (*A, D, G, J*) Raster plot of the neuronal firing. (*B, E, H, K*) The distribution of causal values of each pair of neurons in the whole network. (*C, F, I, L*) ROC curves of the corresponding network with AUC equaling (*C*) 1.0, (*F*) 0.99, (*I*) 1.0, (*L*) 0.98. The parameters are set as (*A-C*) $f = 1.6\,\mathrm{mV}$, $\nu = 0.6\,\mathrm{kHz}$, $S = 0.5\,\mathrm{mV}$, $\Delta t = 0.5\,\mathrm{ms}$, $m = 2$ (time delay is 1 ms), and orders $k = l = 1$, (*D-F*) $f = 2.2\,\mathrm{mV}$, $\nu = 0.3\,\mathrm{kHz}$, $S = 0.6\,\mathrm{mV}$, $\Delta t = 0.5\,\mathrm{ms}$, $m = 6$ (time delay is 3 ms), and orders $k = l = 1$, (*G-I*) $f = 0.5$, $\nu = 0.1\,\mathrm{kHz}$, $S = 0.05$, $\Delta t = 0.5\,\mathrm{ms}$, $m = 6$ (time delay is 3 ms), and orders $k = l = 1$, (*J-L*) $f = 100\,\mu\mathrm{A}\cdot\mathrm{cm}^{-2}$, $\nu = 0.4\,\mathrm{kHz}$, $S = 30\,\mu\mathrm{A}\cdot\mathrm{cm}^{-2}$, $\Delta t = 0.5\,\mathrm{ms}$, $m = 6$ (time delay is 3 ms), and orders $k = l = 1$. The ROC curves for TDCC, TDMI, GC, and TE in (*C, F, I, L*) overlap with each other. The colors are the same as those in Fig. 6.
